# Linking T cell receptor sequence to transcriptional profiles with clonotype neighbor graph analysis (CoNGA)

**DOI:** 10.1101/2020.06.04.134536

**Authors:** Stefan A. Schattgen, Kate Guion, Jeremy Chase Crawford, Aisha Souquette, Alvaro Martinez Barrio, Michael J.T. Stubbington, Paul G. Thomas, Philip Bradley

## Abstract

Multi-modal single-cell technologies capable of simultaneously assaying gene expression and surface phenotype across large numbers of immune cells have described extensive heterogeneity within these complex populations, in healthy and diseased states. In the case of T cells, these technologies have made it possible to profile clonotype, defined by T cell receptor (TCR) sequence, and phenotype, as reflected in gene expression (GEX) profile, surface protein expression, and peptide:MHC (pMHC) binding, across large and diverse cell populations. These rich, high-dimensional datasets have the potential to reveal new relationships between TCR sequence and T cell phenotype that go beyond identification of features shared by clonally related cells. In order to uncover these connections in an unbiased way, we developed a graph-theoretic approach---clonotype neighbor-graph analysis or “CoNGA”---that identifies correlations between GEX profile and TCR sequence through statistical analysis of a pair of T cell similarity graphs, one in which cells are linked based on gene expression similarity and another in which cells are linked by similarity of TCR sequence. Applying CoNGA across diverse human and mouse T cell datasets uncovered known and novel associations between TCR sequence features and cellular phenotype including the classical invariant T cell subsets; a novel defined population of human blood CD8+ T cells expressing the transcription factors *HOBIT* and *HELIOS*, NK-associated receptors, and a biased TCR repertoire, representing a potential previously undescribed lineage of “natural lymphocytes”; a striking association between usage of a specific V-beta gene segment and expression of the *EPHB6* gene that is conserved between mouse and human; and TCR sequence determinants of differentiation in developing thymocytes. As the size and scale of single-cell datasets continue to grow, we expect that CoNGA will prove to be a useful tool for deconvolving complex relationships between TCR sequence and cellular state in single-cell applications.

## Introduction

Previous work pairing gene expression and TCR sequence has largely focused on the TCR sequence as a unique ‘barcode’ by which to identify clonally related cells. Indeed, this approach has revealed important insights into the development and interrelatedness of different T cell subsets within the context of cancer ^1–6^, infectious disease ^7^, and homeostasis ^8^. From these works we see that T cell clones derived from a common clonal ancestor tend to display a similar transcriptional profile. However, the relationship between TCR sequence *similarity* and cellular phenotype has not, to our knowledge, been systematically explored using the large single-cell datasets now available. Researchers have mapped the TCR sequence properties of previously identified T cell subsets ^9–11^, but approaches that can identify completely new populations or subpopulations by correlating GEX and TCR sequence have not been reported. Also lacking are methods for identifying correlations between TCR sequence and GEX that do not extend to global similarity or associate with a defined cell population, for example, correlations between specific TCR sequence properties and expressed genes that might span multiple cell subsets.

In parallel to the developments in single-cell profiling, methods for quantifying TCR repertoire features and identifying patterns within them have matured, helping extend our understanding of T cell biology. Previously, we introduced TCRdist, a measure for assessing inter-TCR similarity capable of identifying closely-related clonotypes based on shared sequence features ^12^. Based on this work and others ^13,14^, it is clear that T cells targeting the same pathogen-derived epitope utilize T cell receptors that share consistent, definable amino acid motifs. In addition to these conventional T cell responses, it is well known that certain unconventional T cell populations, such as mucosal-associated invariant T (MAIT) cells and invariant natural killer T (iNKT) cells, are characterized by conserved TCR sequence features and GEX profiles ^9,10^. The repertoires for a number of distinct T cell subsets with suitable markers for their enrichment have been described, however, it is likely other subsets linked by TCR and GEX remain undiscovered. We hypothesized that by identifying correlations between “TCR neighborhoods”, defined by shared sequence features, and gene expression, we could overcome the strict limitation of examining these correlations within individual clonal families and potentially identify novel associations between T cell antigen-specificities and phenotypes.

To this end, we developed a graph theoretic approach for clonotype neighbor-graph analysis, CoNGA, that identifies correlations between GEX profile and TCR sequence features through analysis of similarity graphs defined on the set of T cell clonotypes and applied it to a collection of publicly-available T cell datasets in an unbiased search for T cell populations linked by covariation in their repertoire features and GEX profiles. In addition to capturing the MAIT and iNKT populations as expected, CoNGA also identified T cell populations for which the linkage was more subtle. These included a *ZNF683+/IKZF2+* (aka *HOBIT*+/*HELIOS*+) population with long and biased CDR3 regions that we hypothesize may represent an unconventional T cell population; CD4 and CD8-positive T cell clusters in mixed PBMC datasets with TCR sequence features that bias CD4 vs CD8 compartment choice; epitope-specific T cell populations; and multiple correlations between gene expression and TCR sequence in a recently published dataset of thymic T cells. Additionally, CoNGA uncovered a striking correlation between expression of the gene *EPHB6*, which flanks the TCR beta locus, and usage of a specific TCR V gene segment, *TRBV30* (*Ephb6* and *TRBV31* in mice). Applying CoNGA to four datasets that included pMHC binding profiles derived from sequencing of cell-surface bound, DNA-barcoded pMHC multimers revealed strong correlations between pMHC binding and both TCR sequence and gene expression. T cell populations specific for individual pMHC epitopes showed distinct gene expression profiles, with EBV epitope-specific T cell populations appearing to cluster according to the stage (latent vs early) of the antigen from which the peptide epitope was derived.

We are not the first to analyze single-cell datasets with parallel TCR and GEX information, however, much of this prior work has used the TCR sequence primarily as a unique tag to identify and track clones. The main contribution of this study is in laying out a systematic approach for discovering relationships between TCR sequence and T cell phenotype in large and heterogeneous single-cell datasets. CoNGA does not require prior identification or isolation of specific subsets in order to identify defining sequence features. CoNGA can also identify GEX/TCR correlations that span multiple T cell clusters rather than simply focusing on one cluster at a time. Thus we are optimistic that as the throughput of single-cell experiments continues to increase, and the dimensionality and multi-modal nature of these experiments continues to grow, graph-based approaches like the one introduced here will play an important role as we leverage these technologies to better understand the adaptive immune system.

## Results

### CoNGA algorithm

CoNGA was developed to identify correlations between gene expression profile and TCR sequence in diverse T cell populations without prior knowledge of the precise nature of these correlations. We envisioned two broad categories of correlation: one based on similarity, in which cells similar with respect to GEX are also similar with respect to TCR sequence, and one based on features, in which specific aspects of GEX and of TCR sequence are correlated, without global similarity of both properties. CoNGA *graph-vs-graph* correlation (details below) was developed to detect the first category of correlation, using the mathematical concept of graph neighborhoods to formalize our intuitive notion of global similarity. De novo discovery of feature-based correlations, without prior knowledge of the correlated features, is more challenging, as it requires enumeration and testing of all possible feature pairs. CoNGA *graph-vs-feature* analysis represents a compromise approach in which we assume that, at least on one side of the correlation, some degree of global similarity is present (this is the “graph-” side); we then enumerate possible features defined by the other property, and test for graph neighborhoods with biased feature distributions. In practice, we find substantial overlap between the results of these two approaches, as, for example, when the identified features in graph-vs-feature correlations are marker genes for a subpopulation of cells that also share detectable global similarity of gene expression. However, we also see cases in which graph-vs-feature analysis reveals a correlation, for example between expression of a specific gene and usage of a particular V gene segment, that is not characterized by global similarity with respect to both gene expression and TCR sequence. These two approaches are also quite complementary: retrospective analysis of graph- vs-graph correlations can, as in the case of the putative MHC-independent population described below, suggest specific gene expression or TCR sequence features that can then be input to graph-vs-feature analysis for sensitive detection of specific correlations.

CoNGA similarity graphs are defined at the level of clonotypes rather than individual cells. We and others have observed that T cells of the same clonotype, which by definition have the same TCR sequence, tend to have similar GEX profiles (**Fig. S1**). Thus, similarity graphs based on gene expression drawn at the level of individual cells will contain many edges connecting cells within the same clonal family. To identify correlations between TCR sequence and gene expression profile beyond the level of individual clonal families, we chose to define similarity graphs at the level of *clonotypes* rather than individual *cells*. Henceforth, for brevity, the term “clonotype” refers to a group of individual cells inferred to be descended from a common clonal ancestor due to their shared expression of a unique, rearranged TCR sequence. In the TCR similarity graph, each node (clonotype) is connected by edges to its K nearest-neighbor (KNN) nodes based on TCR similarity as assessed by the TCRdist measure ^12^, which scores sequence similarity in the pMHC-contacting CDR loops of the TCR alpha and beta chains (here K is an adjustable parameter specified as a fraction of the total number of clonotypes). In the gene expression (GEX) similarity graph, each clonotype is connected by edges to its KNN clonotypes based on similarity in GEX profile (see Methods). Expanded clones are represented by the GEX profile of a single representative cell, the one with the smallest average distance to the rest of the clonal family.

In *graph-vs-graph* correlation analysis **(Fig. 1a,b)**, CoNGA identifies statistically significant overlap between the GEX similarity graph and the TCR similarity graph. We consider each node (clonotype) in turn, count the overlap between its neighbors in the two graphs (i.e., we count how many other nodes are connected to it by both a TCR-similarity edge and a GEX-similarity edge), and assign a significance score that contrasts this observed overlap to that expected under a simple null model: the *CoNGA score* for this clonotype, equal to the hypergeometric probability of seeing the observed overlap by chance, multiplied by the total number of clonotypes, to adjust for multiple testing. CoNGA scores range from 0 to the number of clonotypes; scores close to 0 are significant, scores around 1 are borderline, and scores above 1 are expected to occur by chance (see Methods).This mode of analysis identifies T cell clonotypes whose neighbors in gene expression space overlap significantly with their neighbors in TCR sequence space. Here, we model the concept of a clonotype’s neighbors in GEX or TCR space using the mathematical concept of a *graph neighborhood*, defined as all the vertices directly connected to one central vertex (the colored points in **Fig. 1b**, for example, or the circled points in **Fig. 1d**). CoNGA’s second mode of analysis, *graph-vs-feature* analysis, was developed to detect GEX/TCR correlation that involves specific gene expression or TCR features rather than overall similarity. This mode of analysis can identify TCR sequence neighborhoods with differentially expressed genes (DEGs), for example, or gene expression neighborhoods with distinctive CDR3 sequence features (length, hydrophobicity, charge, etc). In graph-vs-feature correlation analysis **(Fig. 1c,d)**, CoNGA maps numerical features derived from one property (gene expression or TCR sequence) onto the similarity graph defined by the other property and looks for neighborhoods in the graph with unexpectedly high or low feature distributions. The results of CoNGA analyses are summarized in informative visualizations that condense the high-dimensional complexity of these datasets into interpretable plots and graphs.

**Figure 1:**
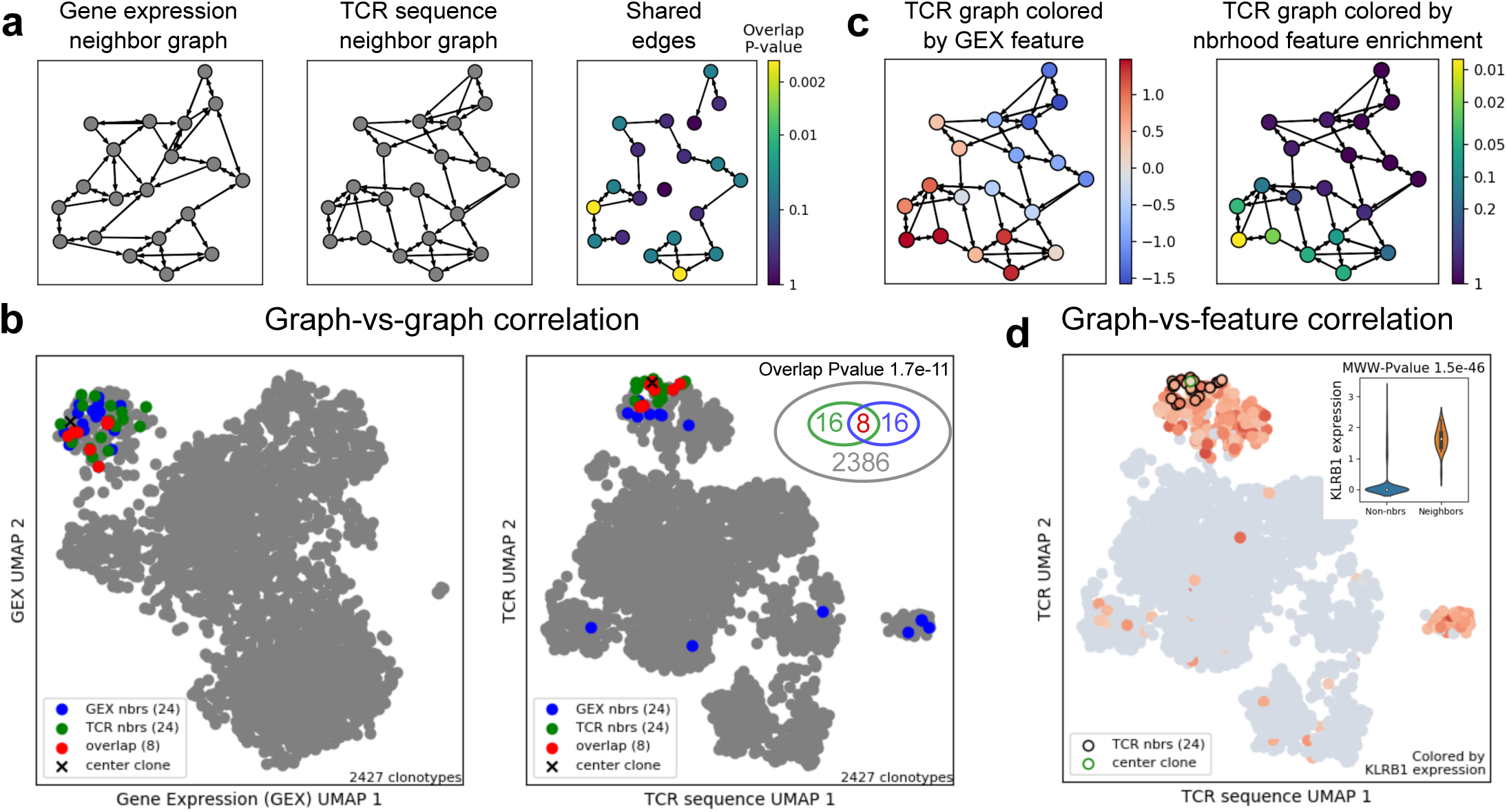
T cell clonotype neighbor graph analysis (CoNGA). **(a)** In graph-vs-graph analysis, CoNGA identifies correlation between T cell gene expression (GEX) and TCR sequence by constructing a gene expression similarity graph and a TCR sequence similarity graph and looking for statistically significant overlap between them. Overlap is assessed on a per-clonotype basis by counting the number of edges that originate at each clonotype and are shared between the two graphs, or equivalently by measuring the overlap between each clonotype’s GEX graph neighbors and its TCR graph neighbors, and assigning a score that reflects the likelihood of seeing equal or greater overlap by chance. **(b)** A single clonotype and its GEX and TCR neighbors are shown in the GEX (left panel) and TCR (right panel) 2D UMAP projections for the *10x_200k_donor2a* dataset. The clonotype is marked with a black ‘x’, its GEX neighbors are shown as blue points, its TCR neighbors as green points, and the clonotypes that are both GEX and TCR neighbors are shown in red. The significance of the observed overlap—8 clones shared between two neighbor sets of size 24 in a total population of 2427 clonotypes—is calculated using the hypergeometric distribution, giving a P value of 1.7 × 10^*−*11^ **(c)** In graph-vs-feature analysis, a numerical feature defined by one property (here gene expression) is mapped onto a similarity graph defined by the other property (TCR sequence), and graph neighborhoods with skewed score distributions are identified using statistical tests that compare the scores for each neighborhood (including the central vertex) with the scores of the remaining clonotypes. **(d)** The gene *KLRB1* (CD161) shows a non-uniform distribution over the TCR sequence landscape—discrete regions of higher expression (red) against a background of lower expression (blue)—suggesting correlation between gene expression and TCR sequence. This is quantified for a single clonotype (green outline) and its TCR sequence neighbors (black outlines) in the inset violin plot, which shows the *KLRB1* expression level for the clonotype and its neighbors on the right and for the remainder of the dataset on the left. The Mann-Whitney-Wilcoxon *P* value for this expression difference is 1.5*×*10^*−*46^

### CoNGA graph-vs-graph analysis identifies correlation between gene expression and TCR sequence

We applied CoNGA to a collection of publicly-available T cell datasets that featured, at a minimum, single-cell GEX and paired TCRαβ sequencing, in an unbiased search for known and novel T cell populations defined by covariation between TCR sequence and GEX profile (see **Table 1** for a list of the datasets analyzed in this work**). Figure 2** illustrates the CoNGA graph-vs-graph analysis workflow for two datasets of human peripheral blood T cells, one a mix of CD4+ and CD8+ cells (*vdj_v1_hs_pbmc*, **Fig. 2a-c**) and one containing flow-sorted CD8+ T cells (*10x_200k_donor2a*, **Fig. 2d-f**; **Table 1**). First, the UMAP algorithm ^15^ is applied to the gene expression and TCRdist matrices of each dataset to generate two dimensional projections of the GEX (**Fig. 2a/d**, left three panels) and TCR landscapes (**Fig. 2a/d**, right three panels). Next, a graph-based clustering algorithm ^16,17^ is applied to the GEX matrix to partition the dataset into clusters of clonotypes with similar transcriptional profiles (**Fig. 2a/d**, panel 1) and to the TCR distance matrix to produce clusters of clonotypes with similar TCR sequences (**Fig. 2a/d**, panel 4). The GEX and TCR landscape projections are colored by CoNGA score to visualize the relative location of the top-scoring CoNGA hits in these landscapes (**Fig. 2a/d**, panels 2 and 5). Finally, the GEX and TCR cluster assignments of CoNGA hits with scores below a threshold (here 1.0) are shown in the 2D projections using bicolored disks whose left (right) half corresponds to the GEX (TCR) cluster assignment (**Fig. 2a/d**, panels 3 and 6 for the GEX and TCR landscapes, respectively).

**Table 1:** Single-cell datasets analyzed in this study

| Dataset | Clonotypes | Cells | CoNGA Hits<sup>a</sup> | Description |
| --- | --- | --- | --- | --- |
| vdj_v1_hs_pbmc | 1539 | 1630 | 113 | CD4 and CD8 T cells from the peripheral blood of a healthy donor<sup>b</sup> |
| vdj_v1_hs_pbmc3 | 2783 | 2945 | 145 | CD4 and CD8 T cells from the peripheral blood of a healthy donor<sup>c</sup> |
| vdj_v1_mm_balbc_pbmc | 1421 | 1423 | 66 | CD4 and CD8 T cells from the peripheral blood of a balbc mouse<sup>d</sup> |
| 10x_200k_donor2a | 2427 | 4721 | 149 | CD8 T cells from the peripheral blood of a healthy donor<sup>e</sup> |
| 10x_200k_donor1 | 20 861 | 33 643 | 1956 | CD8 T cells from the peripheral blood of a healthy donor, sorted for pMHC-multimer binding<sup>f</sup> |
| 10x_200k_donor2 | 8807 | 51 123 | 893 | CD8 T cells from the peripheral blood of a healthy donor, sorted for pMHC-multimer binding<sup>g</sup> |
| 10x_200k_donor3 | 11 971 | 28 748 | 277 | CD8 T cells from the peripheral blood of a healthy donor, sorted for pMHC-multimer binding<sup>h</sup> |
| 10x_200k_donor4 | 10 967 | 20 416 | 125 | CD8 T cells from the peripheral blood of a healthy donor, sorted for pMHC-multimer binding<sup>i</sup> |
| thymus_atlas | 9410 | 9452 | 1044 | Thymic T cells from embryonic, fetal, pediatric, and adult samples<sup>j</sup> |

<sup>a</sup> Graph-vs-graph hits with CoNGA score $\leq 1$
<sup>b</sup> https://support.10xgenomics.com/single-cell-vdj/datasets/2.2.0/vdj_v1_hs_pbmc_5gex
<sup>c</sup> https://support.10xgenomics.com/single-cell-vdj/datasets/3.1.0/vdj_v1_hs_pbmc3
<sup>d</sup> https://support.10xgenomics.com/single-cell-vdj/datasets/3.0.0/vdj_v1_mm_balbc_pbmc_5gex
<sup>e</sup> Ref. 19, https://support.10xgenomics.com/single-cell-vdj/datasets/3.0.2/vdj_v1_hs_aggregated_donor2
<sup>f</sup> Ref. 19, https://support.10xgenomics.com/single-cell-vdj/datasets/3.0.2/vdj_v1_hs_aggregated_donor1
<sup>g</sup> Ref. 19, https://support.10xgenomics.com/single-cell-vdj/datasets/3.0.2/vdj_v1_hs_aggregated_donor2
<sup>h</sup> Ref. 19, https://support.10xgenomics.com/single-cell-vdj/datasets/3.0.2/vdj_v1_hs_aggregated_donor3
<sup>i</sup> Ref. 19, https://support.10xgenomics.com/single-cell-vdj/datasets/3.0.2/vdj_v1_hs_aggregated_donor4
<sup>j</sup> Ref. 27, doi:10.5281/zenodo.3711134

**Figure 2:**
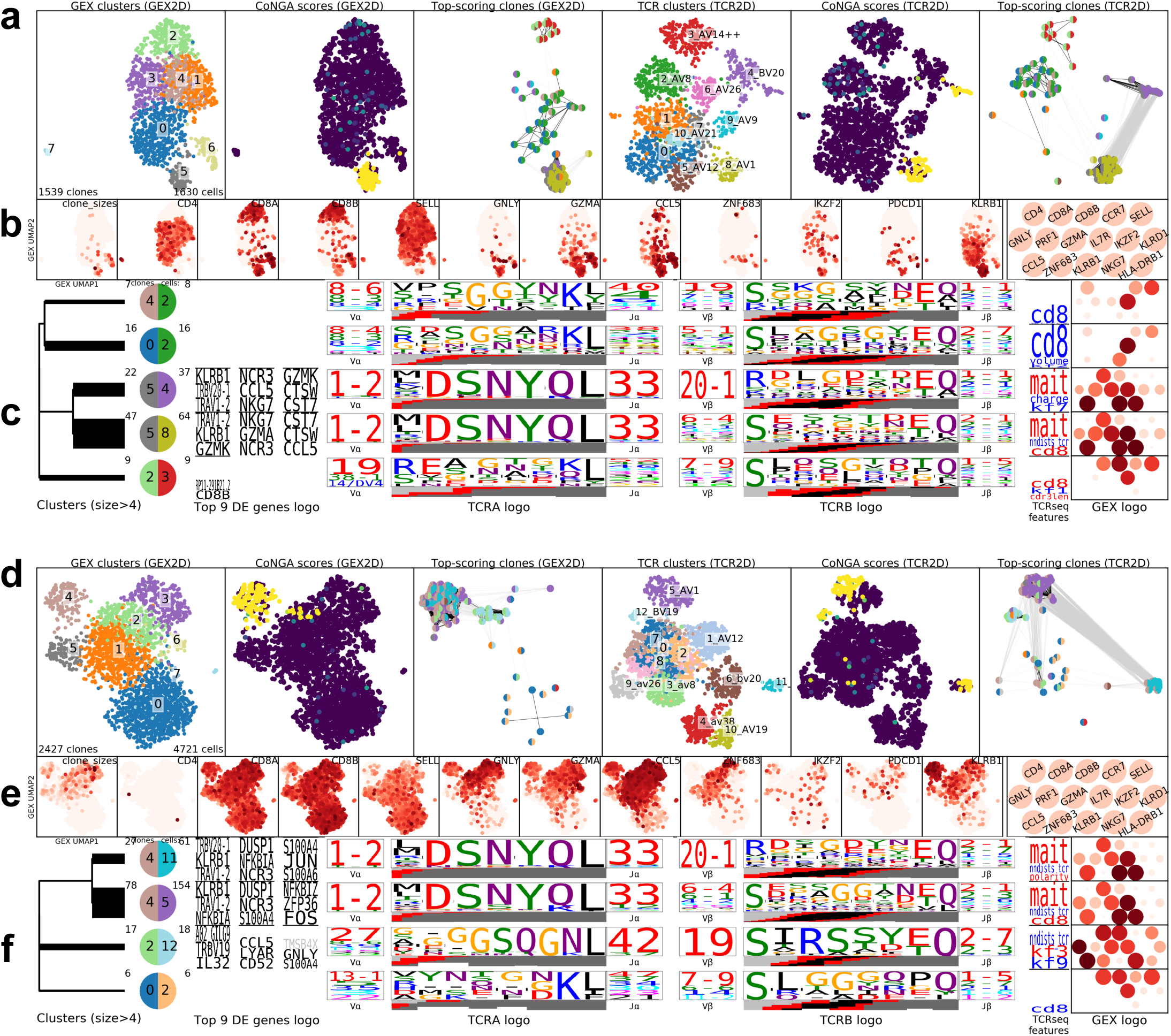
CoNGA identifies GEX/TCR correlation in two datasets of T cells from peripheral blood. **(a-c)** A dataset of mixed CD4+ and CD8+ T cells (*vdj_v1_hs_pbmc*); **(d-f)** a dataset of CD8+ T cells (*10x_200k_donor2a*). **(a,d)** 2D UMAP projections of clonotypes in the dataset based on GEX similarity (left three panels) and TCR similarity (right three panels), colored from left to right by (1) GEX cluster assignment; (2) CoNGA score; (3) GEX/TCR cluster pair assignment, using a bicolored disk whose left half indicates GEX cluster and whose right half indicates TCR cluster (only clones with CoNGA score less than 1 are shown); (4) TCR cluster; (5) CoNGA score; (6) GEX/TCR cluster pair assignment, restricted to clones with CoNGA score less than 1. **(b,e)** Expression of selected marker genes in the GEX UMAP landscape, for visual reference. **(c,f)** Gene expression and TCR sequence features of CoNGA hits in cluster pairs with 5 or more hits are summarized by a series of logo-style visualizations, from left to right: differentially expressed genes (DEGs), TCR sequence logos showing the V and J gene usage and CDR3 sequences for the alpha and beta chains (Dash et al. 2017); biased TCR sequence scores, with red indicating elevated scores and blue indicating decreased scores relative to the rest of the dataset (see **Table S1** for score definitions); expression of a panel of marker genes shown with red disks colored by mean expression and sized according to the fraction of cells expressing the gene (gene names are given above). DEG and TCRseq sequence logos are scaled by the adjusted *P* value of the associations: full logo height requires a top adjusted *P* value below 1*×*10^*−*6^, with partial height and the relative apportionment of height within the logo dictated by the mapping 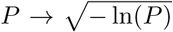. DEGs with fold-change less than 2 are shown in gray. Each cluster pair is indicated by a bicolored disk colored according to GEX cluster (left half) and TCR cluster (right half). The two numbers above each disk show the number of CoNGA hits (on the left) and the total number of cells in those clonotypes (on the right). The dendrogram at the left shows similarity relationships among the cluster pairs based on connections in the GEX and TCR neighbor graphs.

These plots reveal that both datasets contain a substantial number of clonotypes with significant CoNGA scores, and that these CoNGA hits are located in specific regions of the GEX and TCR landscapes. To gain insight into these groups of related clonotypes, we leverage the fact that each dataset has been clustered for both GEX and TCR sequence similarity, independently, and thus each clonotype maps to a pair of clusters (a GEX cluster and a TCR sequence cluster). These cluster pairs provide useful handles by which to identify CoNGA hits because they contain information on GEX and TCR, allowing us to map between the two landscapes (which would require a four-dimensional plot for direct visual correspondence). For example, in **Figure 2a** at the top of the GEX landscape we can see a cluster of CoNGA hits which all belong to GEX cluster 2 (light green on the left half of the disk) and TCR cluster 3 (red on the right half of the disk), or equivalently, cluster pair *(2,3)*; we can infer that these correspond to the group of clonotypes in the TCR landscape also located near the top of the plot, that they are likely CD8+ (from the thumbnail in **Fig. 2b**), and largely TRAV14 (from the TCR cluster identifier in **Fig. 2a)**. Each cluster pair containing an arbitrary minimum number of CoNGA hits (here 5) is characterized by a row of sequence-logo ^18^ style visualizations (**Fig. 2c/f**) that identify the distinguishing features of those CoNGA hits, including the most significant DEGs, TCR gene segment usage, CDR3 motifs, and a GEX logo highlighting several hallmark genes defining canonical T cell subsets (CD4, CD8, etc.). These are arranged in a consistent format that can be scanned for rapid assessment of a cluster’s position within major cell subsets.

Five CoNGA cluster pairs of size 5 or greater were identified in the dataset of mixed CD4 and CD8 human T cells (**Fig. 2c)**. The two largest clusters---*(5,8)* and *(5,4)*, where the first number in each pair indicates the GEX cluster and the second the TCR cluster and we shorten ‘cluster pair’ to ‘cluster’ where context allows---represent MAIT cells: they show high expression of the gene *KLRB1* (CD161) and an invariant TRAV1-2/TRAJ33 alpha chain and restricted Vβ usage. Cluster *(2,3)* contains naive phenotype CD8+ cells that score highly on a sequence-based CD8 compartment preference score (red ‘cd8’ in the ‘TCRseq features’ column; see Methods). Their significant CoNGA scores may reflect the presence of shared sequence features that bias toward the CD8 phenotype and hence correlate with greater similarity of gene expression. Similarly, clusters *(0,2)* and *(4,2)* contain CD4+ cells and score low on the CD8 sequence score (blue ‘cd8’ in the TCRseq features column) and hence may reflect shared TCR sequence features and gene expression consistent with CD4+ fate choice. Application of CoNGA to a second human PBMC dataset and to a mouse PBMC dataset yielded similar results **(Fig. S2)**, with iNKT cells replacing MAIT cells as the dominant invariant subset in the mouse. Turning to the dataset of human CD8+ T cells **(Fig. 2f)**, we again see two MAIT clusters, *(4,11)* and *(4,5)*, differentiated by their TCR beta chain V gene usage (*TRBV20* versus *TRBV6*). Cluster *(2,12)* is characterized by a strong TCR beta chain sequence motif and high expression of cytotoxicity/activation markers including *GNLY* and *CCL5*. The TCR sequence motif matches the consensus for the response to the immunodominant A*02:01-restricted Influenza M1_58_ epitope ^12^. The assignment of this specificity to these cells is supported by the fact that the top DEG for this cluster (‘A02_GILG9’) is actually the read count for a DNA-barcoded A*02:01-M1_58_ multimer that was included in the experiment (note that these pMHC read counts were used for cluster annotation by differential expression analysis but were excluded from the CoNGA neighbor graph construction, 2D projection, and clustering steps so as not to bias the results).

### CoNGA defines a *HOBIT+*/*HELIOS+* T cell population shared across multiple donors

We next applied CoNGA to four large datasets of peripheral blood CD8+ T cells that were enriched for binding to a panel of 50 DNA-barcoded pMHC multimers (*10x_200k_donor1-4* ^19^). The majority of these cells were sorted for positive binding to at least one of the pMHC multimers, and indeed our analysis of TCR:pMHC binding described below finds a number of strong epitope-specific responses. For a few of the pMHC multimers, however, we observed significant levels of non-specific binding **(Fig. S3)**, for example to MAIT cells, or to cells that were very likely part of epitope-specific responses to other epitopes. For this reason, these datasets also include diverse T cells whose binding specificity extends beyond the pMHC multimer panel. CoNGA detected a large number of significant GEX/TCR correlations across these datasets, identifying 62 cluster pairs of size at least 5 **(Figs. S4-7)** and 42 using the more stringent size threshold of .1% of the dataset. **Figure 3** provides an overview of all cluster pairs with at least 21 CoNGA-identified clonotypes (.1% threshold) in the *10x_200k_donor1* dataset. Further examination allowed categorization of the CoNGA cluster pairs depicted in **Figure 3** into three groups: (1) Flu M1_58_-responding clones; (2) MAIT cells; (3) a population of clonotypes with a shared expression profile (high expression of genes including the transcription factors *ZNF683 (*aka *HOBIT) and IKZF2 (*aka *HELIOS)*, along with *DUSP1/2, CD7, CD99*, and *KLRD1*), diverse TCR gene usage, and rather long CDR3 regions.

**Figure 3:**
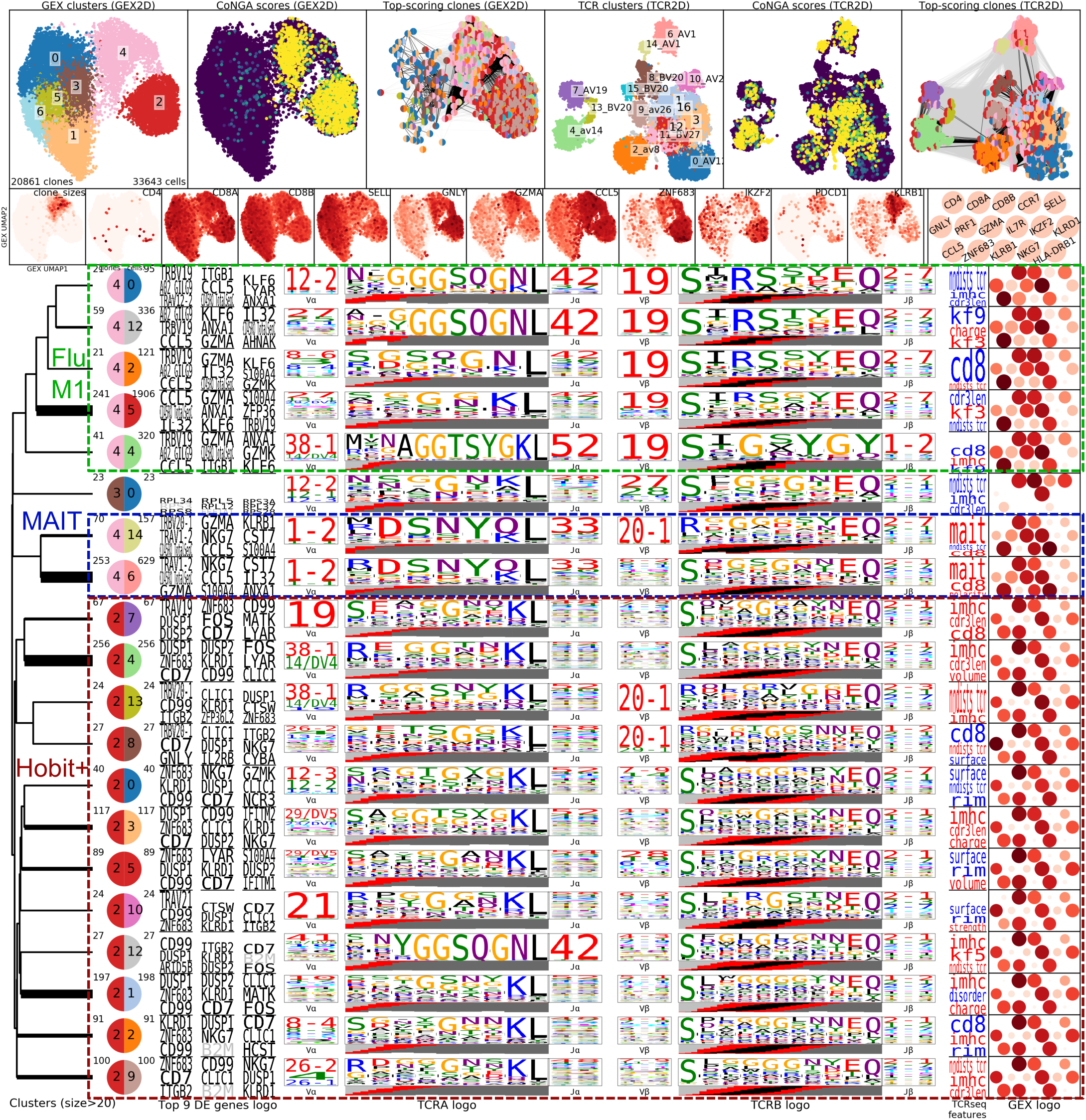
CoNGA plots and cluster logos for a large dataset of CD8+ T cells (*10x_200k_xsdonor1*). Same arrangement of plots as in **Figure 2**. The three colored and dashed boxes group related cluster pairs.

To gain further insight into the large population of *HOBIT*-expressing clonotypes identified by CoNGA, we compared their TCR sequences to a background set formed by pooling all the remaining TCR sequences in the dataset **(Table 2;** see **Table S1** and **Fig. S8** for details on the amino acid property scores). As expected from examination of the TCR sequence logos in **Figure 3**, the CDR3α and CDR3β loops are significantly longer in the *HOBIT*+ CoNGA population than in background (P<10^−300^). The CDR3s are also (1) more positively charged (P<10^−40^); (2) higher in aromatic residues, particularly tryptophan (P<10^−60^), and hydrophobic and bulky amino acids in general (low ‘surface’ and high ‘volume’ scores in **Table 2**); and (3) higher in cysteine (>100-fold enriched in the CDR3β, P<10^−50^). These sequence characteristics are strikingly similar to features identified in a comparison of MHC-independent versus MHC-restricted TCR sequences from an experimental study of TCR repertoires in MHC-knockout mice ^20^. Similar trends were also seen in comparisons of simulated and measured TCR sequences from pre-versus post-selection repertoires ^21–23^, and in CD8aa intraepithelial lymphocytes and their thymic precursors ^24,25^.

**Table 2:**
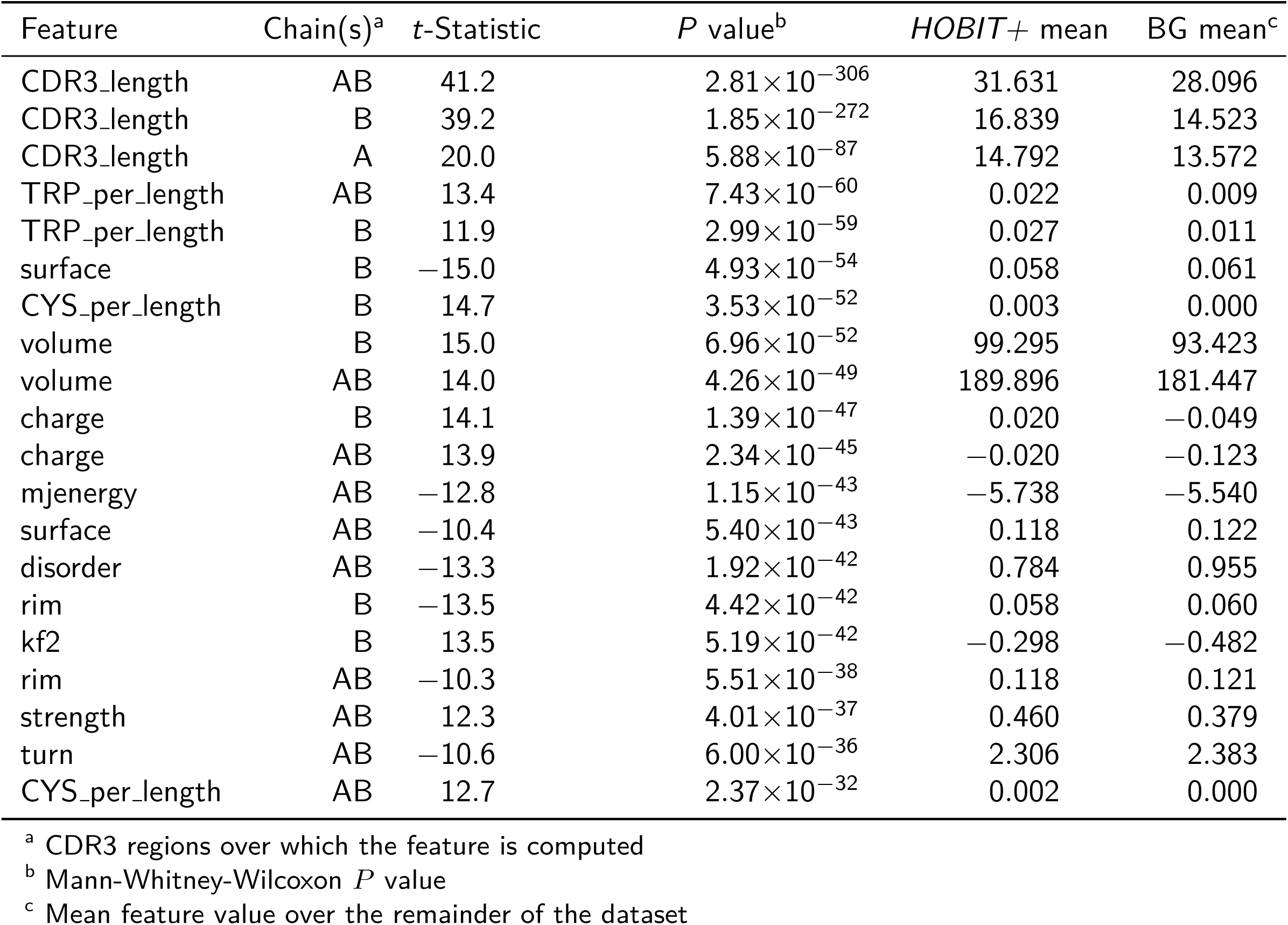
Top 20 sequence features of the *HOBIT+* subset in *10x_200k_donor1*

Depletion of cysteine from the CDR3 loops of MHC-restricted TCRs has been hypothesized to reflect a penalty for disulfide bond formation with cysteines in MHC-presented peptides imposed by negative selection in the thymus; hydrophobic residues positioned within the apex of the CDR3 region are important for mediating interactions with self-peptide MHC in the thymus ^23^. Based on these trends, we hypothesize that this CoNGA-identified population represents a noncanonical, self-specific or MHC-independent, T cell population. To facilitate analysis, we developed a numerical score, the *iMHC score* (for ‘independent of pMHC’), that captures their defining CDR3 sequence features (see Methods and **Table S2**).

We next sought to determine the frequency of the *HOBIT*+ population in peripheral blood T cells based on putative cell surface markers identified from their DEGs. Analysis of the features distinguishing the *HOBIT*+ population in *10x_200k_donor1* suggested that they were likely CD45RA^+^ CD45RO^dim^ based on TotalSeq labeling, negative for *CCR7* expression, and positive for *KLRC2, KLRC3*, and a number of *KIR* genes **(Fig. 4a)**. Therefore, we predicted the *HOBIT*+ cells would be CD45RA^+^ CD45RO^dim/-^ CCR7^-^ KLRC2^+^ KLRC3^+^ KIR^+/-^ in their surface marker phenotype (see **Fig. S9** for gating strategy). We were unable to examine the protein levels of either HOBIT or KLRC3 directly due to the lack of commercially available antibodies. Notably, in the report describing the generation of a HOBIT monoclonal antibody its expression was found to be highest in CD45RA^+^ CCR7^-^ CD8 T cells ^26^. Labeling of PBMC samples from healthy blood donors with these cell surface markers for flow cytometric analysis confirmed the presence of CD45RA^+^ CD45RO^dim/-^ CCR7^-^ CD8 T cells expressing all combinations of KLRC2 and KIR2D (i.e. KLRC2^+^KIR2D^-^, KLRC2^+^KIR2D^+^, and KLRC2^-^KIR2D^+^) **(Fig. 4b)**. The presence of both KLRC2^+^ KIR2D^-^ and KLRC2^+^KIR2D^+^ populations is consistent with the ubiquitous *KLRC2* expression and stochastic KIR2D expression within the *HOBIT*+ population of *10x_200k_donor1.* However, the KLRC2^-^KIR2D^+^ phenotype is inconsistent with these criteria and likely represents a distinct (but sizable) CD8 subset. As a percentage of total PBMC CD8 T cells, the KLRC2^+^ KIR2D^+/-^ subset is in the range of 0.2-10.1% while KLRC2^-^ KIR2D^+^ cells ranged between 0.3-7.6% (n = 11) **(Fig. 4c)**. We next sorted the KLRC2^+^ KIR2D^+/-^ and KLRC2^-^ KIR2D^+^ CD8 T cells and measured *ZNF683, KLRC2*, and *KLRC3* expression within these populations relative to each donors’ own sorted CD8+CD45RA-CD45RO+ memory subset using qRT-PCR. Here, we found expression of *KLRC2* and *KLRC3* was enriched in the KLRC2^+^ KIR2D^+/-^ CD8 T cells, and to a lesser extent in the KLRC2^-^ KIR2D^+^ subset **(Fig. 4d)**. However, *ZNF683* appeared to be enriched only within the KLRC2^+^ KIR2D^+/-^ subset, supporting their identity as the putative *HOBIT*+ population and further suggesting KLRC2^-^ KIR2D^+^ T cells are in fact a separate, distinct subset. Taken together, these data confirm the existence of CD8+ CD45RA^+^ CD45RO^dim/-^ CCR7^-^ KLRC2^+^ KIR2D^+/-^ T cells in peripheral blood expressing *ZNF683* consistent with the *HOBIT*+ population, and that this subset, while variable across individuals, comprises a sizable fraction of the CD8 T cells (up to 10% in some individuals).

**Figure 4:**
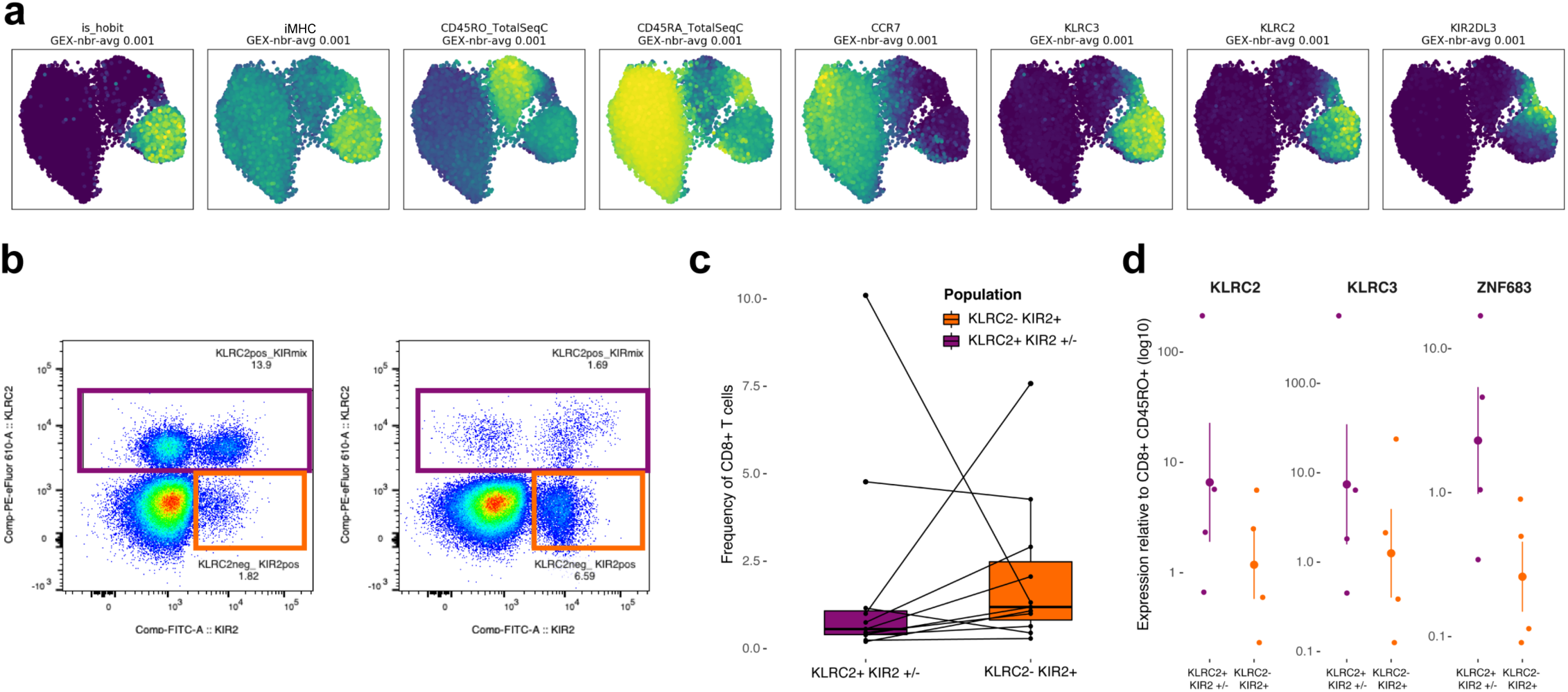
Identification of KLRC2+ CD8 T cells as the *HOBIT+* iMHC-elevated population. **(a)** 2D GEX projection of the *10x_200k_donor1* dataset colored by *‘is_hobit’* (an indicator variable for the *HOBIT+* CoNGA population), iMHC score, CD45RA and CD45RO TotalSeq, *CCR7, KLRC2, KLRC3*, and *KIR2DL3* expression averaged in its GEX graph neighborhood (with neighborhood size equal to 0.1% of the dataset). The *is hobit* variable is 1 for all CoNGA hits in GEX cluster 2 and 0 otherwise. **(b)** Detection of KLRC2+ KIR2D+/- and KLRC2-KIR2D+ CD8 T cells in human PBMCs of two representative donors. Gated on lineage-, CD56+/-, CD3+, CD8+, CD4-, CD45RA+, CD45ROdim/-, CCR7-, CD248-cells (full gating strategy in **Figure S10**). **(c)** Frequency of cell populations in panel (b) as frequency of CD8 T cells. (d) *ZNF683, KLRC2*, and *KLRC3* expression in sorted KLRC2+ KIR2D+/- and KLRC2-KIR2D+ CD8 T cells shown as fold-change relative to CD8+ CD45RO+ T cells within each PBMC donor (n=4).

### CoNGA identifies GEX/TCR correlation in thymic T cells

We next applied CoNGA to a recently published single-cell atlas of human thymic T cells ^27^. This dataset combines thymic tissue from embryonic and fetal stages as well as postnatal thymi from children and adults, totaling over 9400 clonotypes with paired alpha and beta TCR sequences. CoNGA identified a large number of significant hits in this rich and complex dataset, primarily within the DP (double-positive), CD8 single positive (SP), CD4 SP, Treg, and CD8αα^+^ thymic populations **(Fig. 5)**. In TCR sequence space, we see a concentration of hits in the TRAV41 cluster (this TRAV gene is enriched in DP cells), the TRAV1 and TRAV12 clusters (enriched in CD8 cells), and in the TRAV14 cluster (enriched in CD8αα cells) **(Fig. 5)**. The CD8+ cluster pairs identified by CoNGA also showed high CD8 sequence scores and high scores for a measure (‘alphadist’) that reflects the genomic distance between the TRAV and TRAJ gene segments incorporated in a clonotype’s TCR alpha chain. The DP cluster pairs show low alphadist scores, preference for *TRAV41* and other TRAV genes at the 3’ end of the locus, longer CDR3 loops (CDR3 length has been shown to decrease during thymic selection ^21^), and higher scores for the rim, surface, and disorder amino acid properties, which may suggest more polar, less bulky, and less strongly interacting CDR3 regions. Consistent with the findings of Park et al., the CD8αα cluster pairs both show low alphadist scores, however, CoNGA further identified high iMHC scores and longer CDR3 loops as TCR features of these clusters. Interestingly, the CD8αα(II) cluster pair expressed both *ZNF683* and *IKZF2*, which together with TCR features similar to those of the *HOBIT*+ T cells in the blood identified above, suggests a possible precursor relationship between these two populations that warrants further investigation.

**Figure 5:**
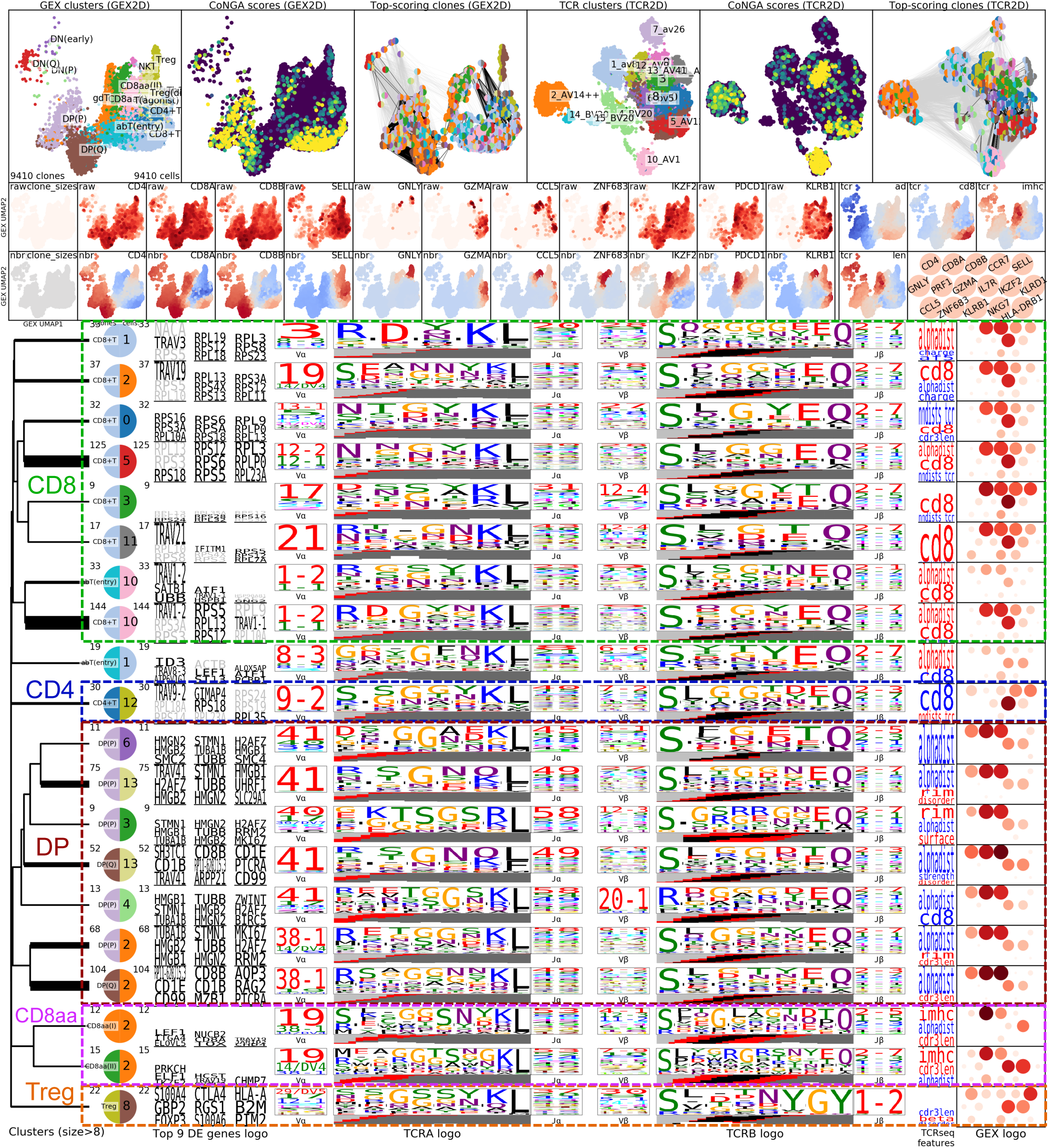
CoNGA plots and cluster logos for a large dataset of thymic T cells (*thymus_atlas*). Same arrangement of plots as in **Figure 2**, with two additions: below the raw GEX landscape thumbnails are added GEX-neighborhood averages of Z-score normalized expression levels, to aid in detecting differential expression of marker genes; to the right of the GEX landscape thumbnails are plotted four TCR feature scores, also GEX-neighborhood averaged (‘ad’ is short for the alphadist score and ‘len’ is short for cdr3len; see main text and **Table S1**). The five colored and dashed boxes group related cluster pairs as annotated by the text labels.

### CoNGA graph-vs-feature analysis confirms sharing of the *HOBIT*+/*HELIOS*+ T cell subset across donors

We have seen that CoNGA graph-vs-graph analysis can identify a variety of correlations between gene expression and TCR sequence, ranging from the invariant MAIT and iNKT lineages, to sequence motifs and expression biases in an epitope-specific response, to the weaker CDR3 sequence preferences and differentially expressed genes that characterize the *HOBIT*+ population (which would likely be difficult to identify from analysis of TCR sequence or gene expression alone). To be detected, these correlations must be characterized by some degree of elevated global similarity in both transcriptional profile and TCR sequence within the relevant cell population. Thus, correlations that involve only a few genes or very specific TCR sequence features, or ones that are not well captured by our global GEX and TCR distance measures, may go undetected. CoNGA graph-vs-feature analysis was developed as a complementary graph-based approach that could detect GEX/TCR correlations that are not characterized by global similarity of both properties. In graph-vs-feature analysis, numerical features calculated on the basis of one cellular property, GEX or TCR sequence, are mapped onto a similarity graph defined by the other property, and the feature score distributions for each of the neighborhoods in the graph are compared to the background distributions to identify neighborhoods with skewed scores (here a graph neighborhood consists of a single central vertex together with all of its directly connected neighbors). As GEX features, we consider the expression levels of individual genes, and for TCR sequence features, we use a set of CDR3 amino acid property values as well as a handful of additional, sequence-based scores (**Table S1** and **Fig. S8**).

We used graph-vs-feature analysis to identify additional members of the *HOBIT*+/*HELIOS*+ unconventional T cell subset by looking for GEX graph neighborhoods with elevated iMHC scores. Although the per-clonotype iMHC score is highly variable **(Fig. 6a)**, by computing averages over GEX graph neighborhoods we can identify a subregion of GEX space with enhanced scores **(Fig. 6b)**, whose significance can be assessed using standard statistical tests **(Fig. 6c)**. Three of the four *10x_200k* donors show populations of clonotypes with significantly enhanced iMHC scores **(Fig. 6c-f)** whose DEGs correlate well with one another and with the key marker genes (*ZNF683, CD7, CD99, DUSP1/2*) for the original *HOBIT*+ CoNGA clusters. Interestingly, the outlier donor with very few iMHC-high clones was also significantly older than the other 3 donors (age 50 versus ages 30, 31, and 38), consistent with an age-related decline of this putative natural T cell population. Comparison of iMHC score distributions for the *HOBIT*+ CoNGA clonotypes to those of TCRs with known MHC restriction **(Fig. S10)** suggests possible affinity with other MHC-independent T cell subsets.

**Figure 6:**
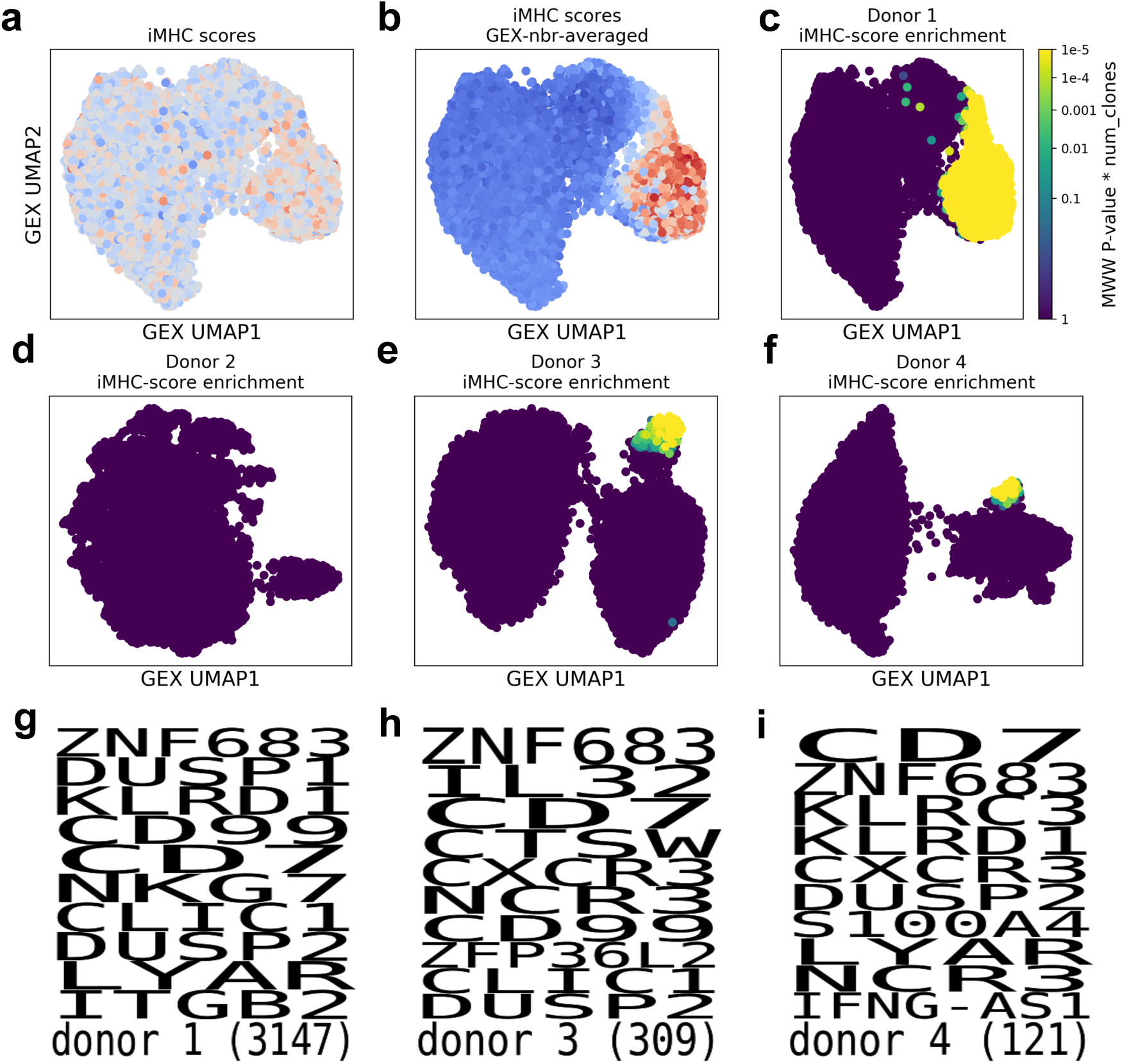
Graph-vs-feature correlation analysis reveals GEX neighborhoods with elevated iMHC scores across multiple donors. **(a)** 2D GEX projection of the *10x_200k_donor1* dataset colored by iMHC score (standardized to have mean 0 and standard deviation 1). **(b)** The same projection as in (a) but each clonotype is colored by the average iMHC score in its GEX graph neighborhood. **(c)** The same projection as in (a) but colored by *P* values for iMHC enrichment in each clonotype’s graph neighborhood (the set of iMHC scores in each clonotype’s neighborhood are compared to the remainder of the iMHC scores using an unpaired, 1-sided Mann-Whitney-Wilcoxon test). **(d)** 2D GEX projection of the *10x_200k_donor2* dataset colored by iMHC score neighborhood enrichment P-values. **(e)** 2D GEX projection of the *10x_200k_donor3* dataset colored by iMHC score neighborhood enrichment P-values. **(f)** 2D GEX projection of the *10x_200k_donor4* dataset colored by iMHC score neighborhood enrichment P-values. **(g)** Top 10 DEGs for the clonotypes with significant iMHC enrichment in the *10x_200k_donor1* dataset. **(h)** Top 10 DEGs for the clonotypes with significant iMHC enrichment in the *10x_200k_donor3 dataset*. **(i)** Top 10 DEGs for the clonotypes with significant iMHC enrichment in the *10x_200k_donor4* dataset. (There were too few clonotypes with significant iMHC enrichment in the *10x_200k_donor2* dataset to identify differentially expressed genes).

### Graph-vs-feature analysis reveals differential gene expression across the TCR landscape

We applied graph-vs-feature analysis in the reverse direction (i.e. between the TCR graph and GEX features) to identify genes that are differentially expressed in specific TCR graph neighborhoods. **Table 3** provides the top hits from this analysis for the datasets analyzed in this study (the top significant gene for each cluster pair and a maximum of 10 genes per dataset are shown). Notable features include MAIT-associated genes such as *KLRB1* and *SLC4A10*; genes associated with the iMHC population such as *ZNF683* and *KLRC3*; and genes upregulated in the M1_58_ response including *ITGB1* and *KLRC1* (in donor 2). We also observed TCR neighborhoods with elevated levels of *CD8A* and *CD8B*, which appear to overlap with the populations identified in the earlier graph-vs-graph correlation analysis and suggest the presence of TCR sequence features that bias toward the CD8+ compartment. Some associations are a consequence of the V(D)J recombination process itself, such as the positive association between TCR neighborhoods using *TRBJ1* family genes and the *TRBC1* constant region, which is deleted during D-J rearrangement involving *TRBJ2*-family genes and hence cannot be used in *TRBJ2*-containing TCRs.

**Table 3:**
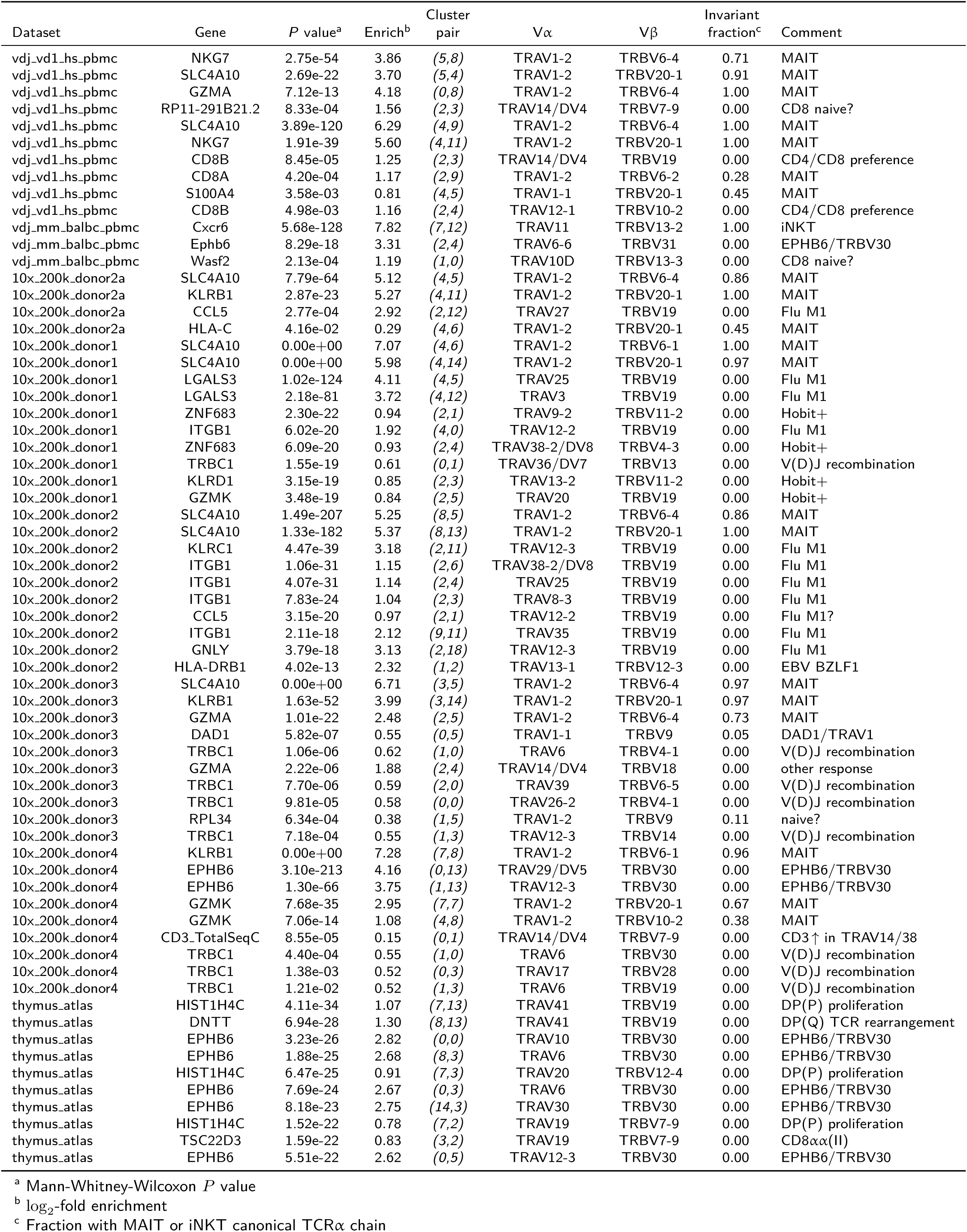
Top differentially-expressed genes in TCR graph neighborhoods

**Figure 7** illustrates four graph-vs-feature correlations, showing visually how specific TCR-based and GEX-based features correlate across the 2D clonotype landscapes. In donor 1 (**Fig. 7a**), the correlation between the iMHC score, a TCR feature, and two GEX features, expression of the genes *ZNF683* and *KLRC3*, is shown by coloring the clonotypes in the GEX UMAP projection by these three features, averaged over GEX graph neighborhoods. Here, the averaging serves to reduce noise and also to highlight TCR feature trends that are consistent with the GEX similarity structure (since we are averaging over the GEX graph neighborhoods). In donor 2 (**Fig. 7b**), we can see correlation, now over the TCR landscape and averaged over TCR graph neighborhoods, between a TCR feature, cell-surface bound A*02:01-M1_58_ pMHC, and expression levels of two genes that mark the M1-responding clonotypes. In donor 3 (**Fig. 7c**), we see correlation over the TCR landscape between a TCR feature, occurrence of the canonical MAIT alpha chain, and the expression of two MAIT cell marker genes. Finally, in donor 4 (**Fig. 7d**), we can see the correlation over the TCR landscape between a TCR feature, usage of the *TRBV30* gene segment, and expression of the gene *EPHB6*.

**Figure 7:**
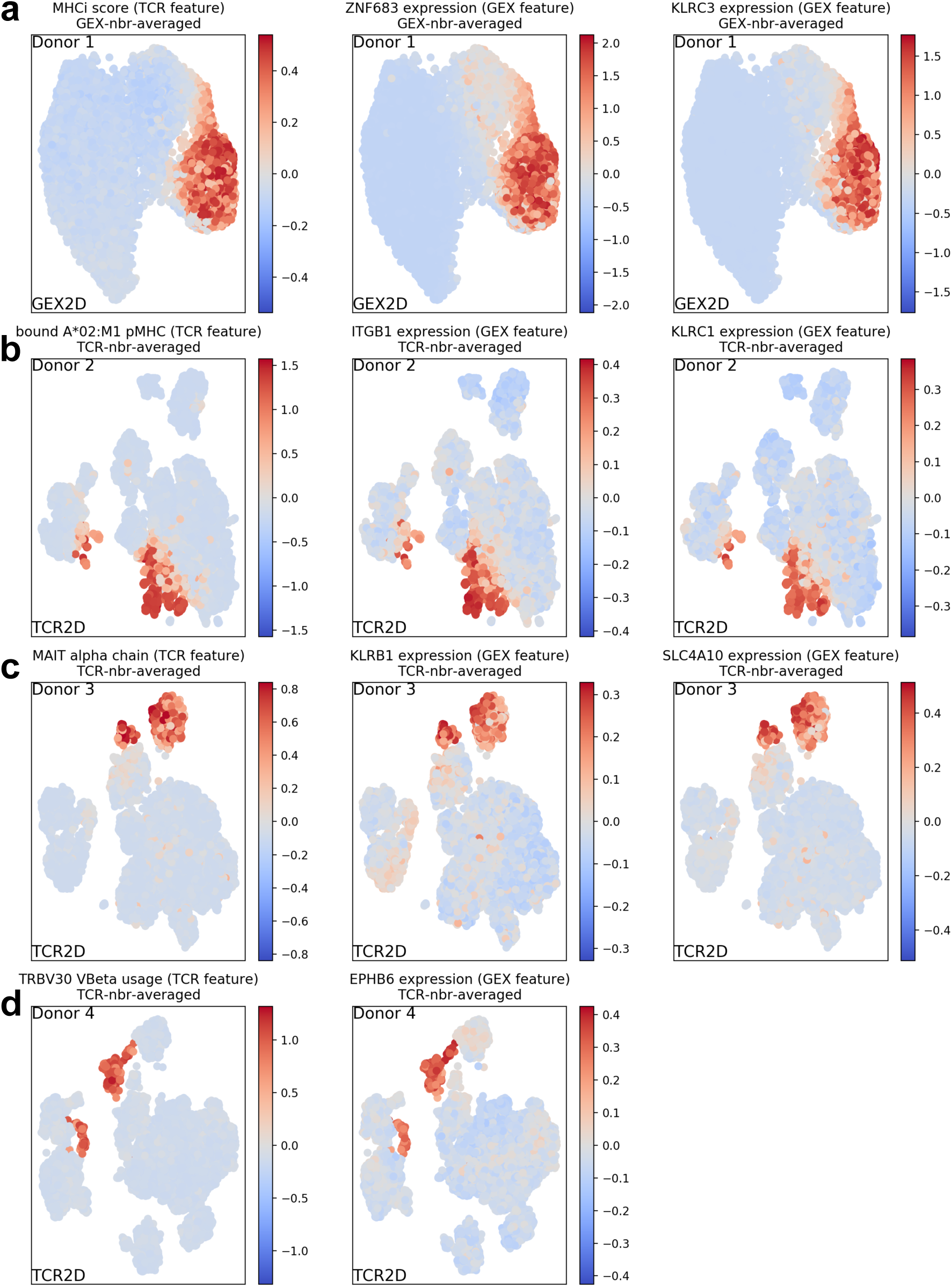
Graph-vs-feature correlation analysis highlights TCR:GEX covariation. In each of the four rows, correlation between a score derived from the TCR sequence (left panel) and 1-2 scores derived from the GEX profile (right panels) is illustrated by mapping the scores onto the 2D UMAP GEX or TCR landscape for the given dataset (after Z-score normalizing and averaging over graph neighborhoods). **(a)** iMHC score averaged over GEX neighborhoods correlates well with marker genes *ZNF683* and *KLRC3* for the HOBIT+ population. **(b)** *KLRB1* and *SLC4A10* averaged over TCR neighborhoods correlate with the MAIT cell population as defined by TCR*α* sequence. **(c)** *ITGB1* and *KLRC1* are elevated in the Flu A*02:M1_58_ response, here defined by the surface counts for the multimerized A*02:M1_58_ pMHC. **(d)** T cells with TRBV30-containing TCRs have elevated *EPHB6* expression.

Our TCR graph-based differential expression analysis identified several associations with the *EPHB6* gene (and its murine homolog *Ephb6)*, which codes for the Ephrin-B receptor Type 6 protein EPHB6 (**Table 3**, for example, the top non-MAIT association for *10x_200k_donor4* and the top non-iNKT association for *vdj_v1_mm_balbc_pbmc*). A recurring feature of these associations is the usage of the *TRBV30* gene segment (*TRBV31* in mouse). A focused search for covariation between TCR gene segment usage and gene expression using differential expression analysis confirmed a strong tendency for higher *EPHB6* expression in clonotypes that incorporate the *TRBV30* gene segment (or *TRBV31* in mouse; **Fig. 8 and Table S3**). The *TRBV30* segment is unique among TRBV genes in being located downstream of the TRBJ and TRBC genes at the end of the TCR beta locus; incorporation of TRBV30 into the TCR by V(D)J recombination requires an altered joining process in which intervening DNA sequence is inverted rather than being deleted ^28^. Providing a potential clue into the mechanism underlying this covariation, *EPHB6* is located adjacent to *TRBV30* on Chromosome 7, ∼40kb downstream from the TCR beta locus **(Fig. 8a)**. The strong correlation between *TRBV30* usage and *EPHB6* expression may indicate that expression of a *TRBV30*-containing TCR transcript also boosts expression of the *EPHB6* gene (the mouse *TRBV31* gene segment is located at an analogous location to that of *TRBV30* in the mouse TR locus, and is also directly adjacent to the mouse homolog *Ephb6*). Given that EPHB6 has been shown to play a role in T cell activation ^29,30^, *TRBV30*+ clonotypes may have distinctive functional properties due to their elevated expression of the *EPHB6* transcript. We also observed covariation, albeit weaker, between *TRAV1-1* usage and expression of the *DAD1* (Defender against cell death 1) gene **(Table S3)**, which flanks the TCR alpha locus at a position analogous to that of *EPHB6*. Given that *TRAV1-1* and *DAD1* are located at opposite ends of the TCR alpha locus, the mechanism underlying this correlation is less clear. Together, these findings show an interaction between the usage of TCR genes at the edges of the TCR loci and the expression of non-TCR genes flanking the loci.

**Figure 8:**
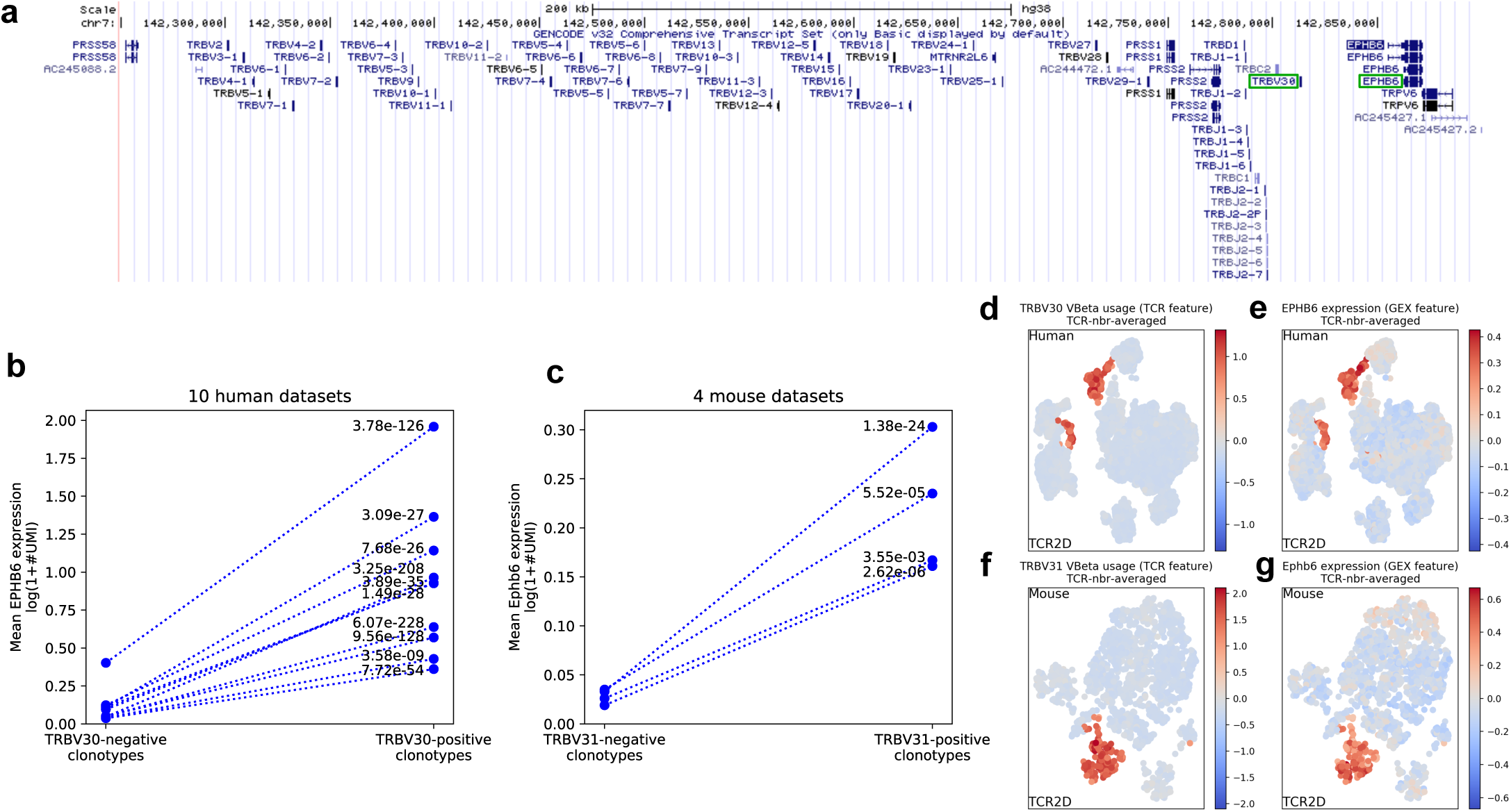
TRBV30 gene usage and *EPHB6* expression are correlated. **(a)** Genomic locations of TRBV30 and *EPHB6* at the 3’ end of the human TCRβ locus. **(b-c)** Average *EPHB6* expression for TRBV30-negative and TRBV30-positive clonotypes in (b) 9 human datasets and (c) 3 mouse datasets. Each marker represents the average over all (TRBV30*−* or TRBV30+) clonotypes for a single dataset. The two markers for each dataset are connected by a dashed line. **(d-g)** 2D projections based on TCR sequence of a mouse (d-e) and human (f-g) dataset colored by TRBV30 (TRBV31 in mouse) usage (d,f) and *EPHB6* expression (e,g) averaged over TCR neighborhoods. Strong correlation is evident between the two features, one derived from the TCR sequence and one from the gene expression profile.

### Neighbor-graph analysis of TCR:pMHC binding highlights GEX similarity among T cells that recognize the same epitope

The use of pMHC-multimers conjugated to DNA barcodes as cell labeling reagents enables high-throughput interrogation of pMHC binding in parallel with other single-cell analyses. We applied CoNGA to investigate correlation between gene expression profiles, TCR sequences, and pMHC:TCR interactions in a large dataset of human T cells sorted for pMHC-multimer binding (*10x_200k_donor1-4*). To do this, we used the pMHC-binding information, stringently filtered and condensed to the level of clonotypes (see Methods), to define a new neighbor graph structure in which edges connect clones that bind to the same pMHC. We then applied CoNGA graph-vs-graph analysis to look for statistically significant overlap between this pMHC-binding graph and the GEX and TCR similarity graphs defined above. We measured graph overlap, on a per-pMHC basis, as the enrichment of GEX (or TCR) similarity graph edges within the pMHC positive clonotypes. Specifically, for each pMHC, we looked to see whether there were more GEX (or TCR) similarity edges within the set of clonotypes positive for that pMHC than we would expect by chance, and quantified this graph overlap by computing a fold-enrichment as well as an approximate P-value **(Table 4, Figure 9)**. From this analysis we can see, as expected, that nearly all the pMHC-positive clonotype subsets show greater than expected TCR sequence similarity. Indeed, the only pMHCs with a negative TCR neighbor-enrichment score are A03_KLG, which appears to show high levels of non-specific binding (**Fig. S3**), and B08_RAK in donor 1, who is HLA-B*08:01 negative. Moreover, pMHCs with large numbers of analyzed clonotypes show highly significant TCR similarity as assessed by the TCR-pMHC graph overlap. Interestingly, we also see that all pMHC-positive populations show greater than expected GEX similarity, with highly significant P-values and large fold-enrichments for most pMHCs with a sufficient number of analyzed clones. These results suggest that clonotypes positive for the same pMHC have more similar gene expression profiles than would be expected by chance.

**Table 4:**
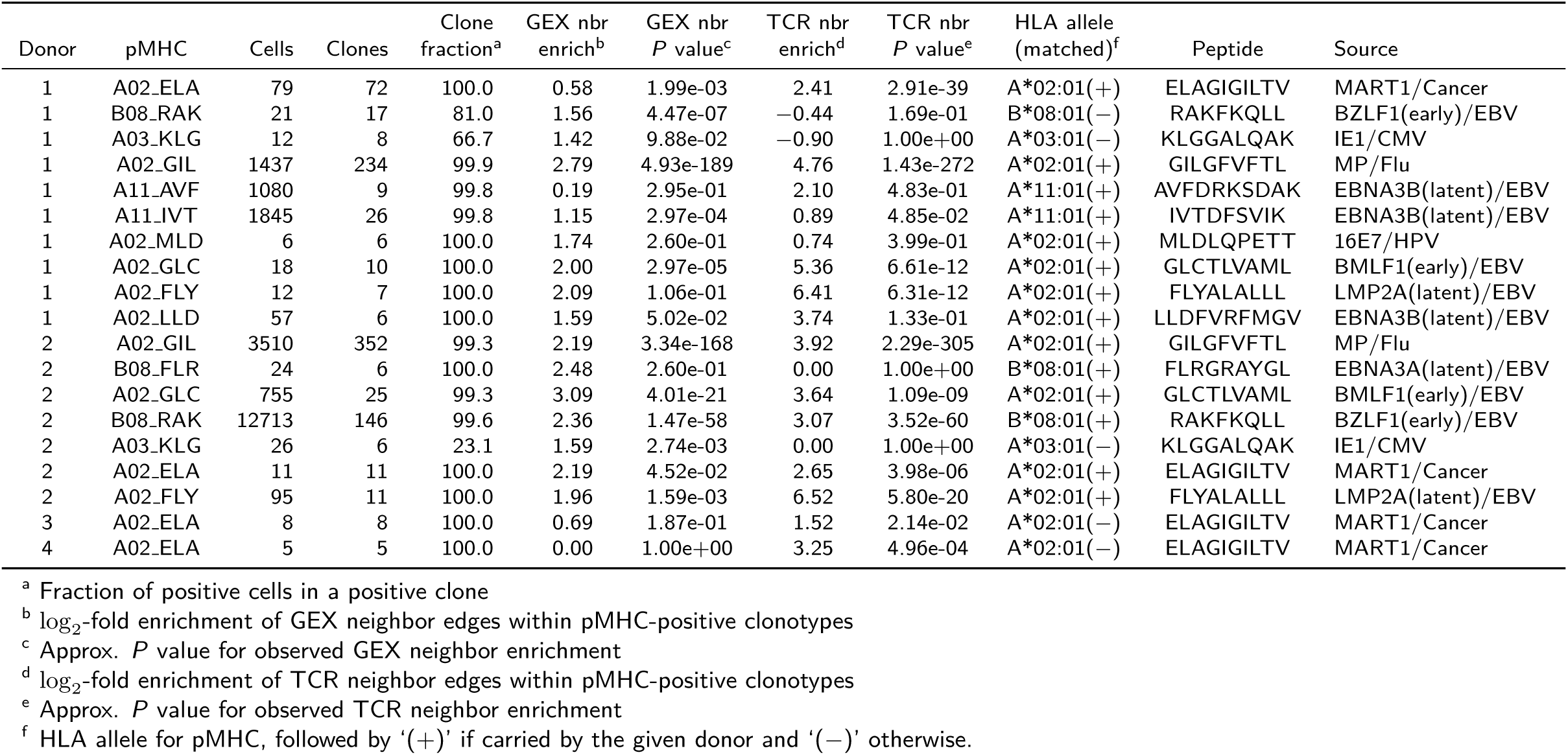
pMHC binding analysis

| Donor | pMHC | Cells | Clones | Clone fraction<sup>a</sup> | GEX nbr enrich<sup>b</sup> | GEX nbr *P* value<sup>c</sup> | TCR nbr enrich<sup>d</sup> | TCR nbr *P* value<sup>e</sup> | HLA allele (matched)<sup>f</sup> | Peptide | Source |
| --- | --- | --- | --- | --- | --- | --- | --- | --- | --- | --- | --- |
| 1 | A02.ELA | 79 | 72 | 100.0 | 0.58 | 1.99e-03 | 2.41 | 2.91e-39 | A*02:01(+) | ELAGIGILTV | MART1/Cancer |
| 1 | B08.RAK | 21 | 17 | 81.0 | 1.56 | 4.47e-07 | -0.44 | 1.69e-01 | B*08:01(-) | RAKFKQLL | BZLF1(early)/EBV |
| 1 | A03.KLG | 12 | 8 | 66.7 | 1.42 | 9.88e-02 | -0.90 | 1.00e+00 | A*03:01(-) | KLGGALQAK | IE1/CMV |
| 1 | A02.GIL | 1437 | 234 | 99.9 | 2.79 | 4.93e-189 | 4.76 | 1.43e-272 | A*02:01(+) | GILGFVFTL | MP/Flu |
| 1 | A11.AVF | 1080 | 9 | 99.8 | 0.19 | 2.95e-01 | 2.10 | 4.83e-01 | A*11:01(+) | AVFDRKSDAK | EBNA3B(latent)/EBV |
| 1 | A11.IVT | 1845 | 26 | 99.8 | 1.15 | 2.97e-04 | 0.89 | 4.85e-02 | A*11:01(+) | IVTDFSVIK | EBNA3B(latent)/EBV |
| 1 | A02.MLD | 6 | 6 | 100.0 | 1.74 | 2.60e-01 | 0.74 | 3.99e-01 | A*02:01(+) | MLDLQPETT | 16E7/HPV |
| 1 | A02.GLC | 18 | 10 | 100.0 | 2.00 | 2.97e-05 | 5.36 | 6.61e-12 | A*02:01(+) | GLCTLVAML | BMLF1(early)/EBV |
| 1 | A02.FLY | 12 | 7 | 100.0 | 2.09 | 1.06e-01 | 6.41 | 6.31e-12 | A*02:01(+) | FLYALALLL | LMP2A(latent)/EBV |
| 1 | A02.LLD | 57 | 6 | 100.0 | 1.59 | 5.02e-02 | 3.74 | 1.33e-01 | A*02:01(+) | LLDFVRFMGV | EBNA3B(latent)/EBV |
| 2 | A02.GIL | 3510 | 352 | 99.3 | 2.19 | 3.34e-168 | 3.92 | 2.29e-305 | A*02:01(+) | GILGFVFTL | MP/Flu |
| 2 | B08.FLR | 24 | 6 | 100.0 | 2.48 | 2.60e-01 | 0.00 | 1.00e+00 | B*08:01(+) | FLRGRAYGL | EBNA3A(latent)/EBV |
| 2 | A02.GLC | 755 | 25 | 99.3 | 3.09 | 4.01e-21 | 3.64 | 1.09e-09 | A*02:01(+) | GLCTLVAML | BMLF1(early)/EBV |
| 2 | B08.RAK | 12713 | 146 | 99.6 | 2.36 | 1.47e-58 | 3.07 | 3.52e-60 | B*08:01(+) | RAKFKQLL | BZLF1(early)/EBV |
| 2 | A03.KLG | 26 | 6 | 23.1 | 1.59 | 2.74e-03 | 0.00 | 1.00e+00 | A*03:01(-) | KLGGALQAK | IE1/CMV |
| 2 | A02.ELA | 11 | 11 | 100.0 | 2.19 | 4.52e-02 | 2.65 | 3.98e-06 | A*02:01(+) | ELAGIGILTV | MART1/Cancer |
| 2 | A02.FLY | 95 | 11 | 100.0 | 1.96 | 1.59e-03 | 6.52 | 5.80e-20 | A*02:01(+) | FLYALALLL | LMP2A(latent)/EBV |
| 3 | A02.ELA | 8 | 8 | 100.0 | 0.69 | 1.87e-01 | 1.52 | 2.14e-02 | A*02:01(-) | ELAGIGILTV | MART1/Cancer |
| 4 | A02.ELA | 5 | 5 | 100.0 | 0.00 | 1.00e+00 | 3.25 | 4.96e-04 | A*02:01(-) | ELAGIGILTV | MART1/Cancer |

<sup>a</sup> Fraction of positive cells in a positive clone
<sup>b</sup> log<sub>2</sub>-fold enrichment of GEX neighbor edges within pMHC-positive clonotypes
<sup>c</sup> Approx. *P* value for observed GEX neighbor enrichment
<sup>d</sup> log<sub>2</sub>-fold enrichment of TCR neighbor edges within pMHC-positive clonotypes
<sup>e</sup> Approx. *P* value for observed TCR neighbor enrichment
<sup>f</sup> HLA allele for pMHC, followed by '(+)' if carried by the given donor and '(-)' otherwise.

**Figure 9:**
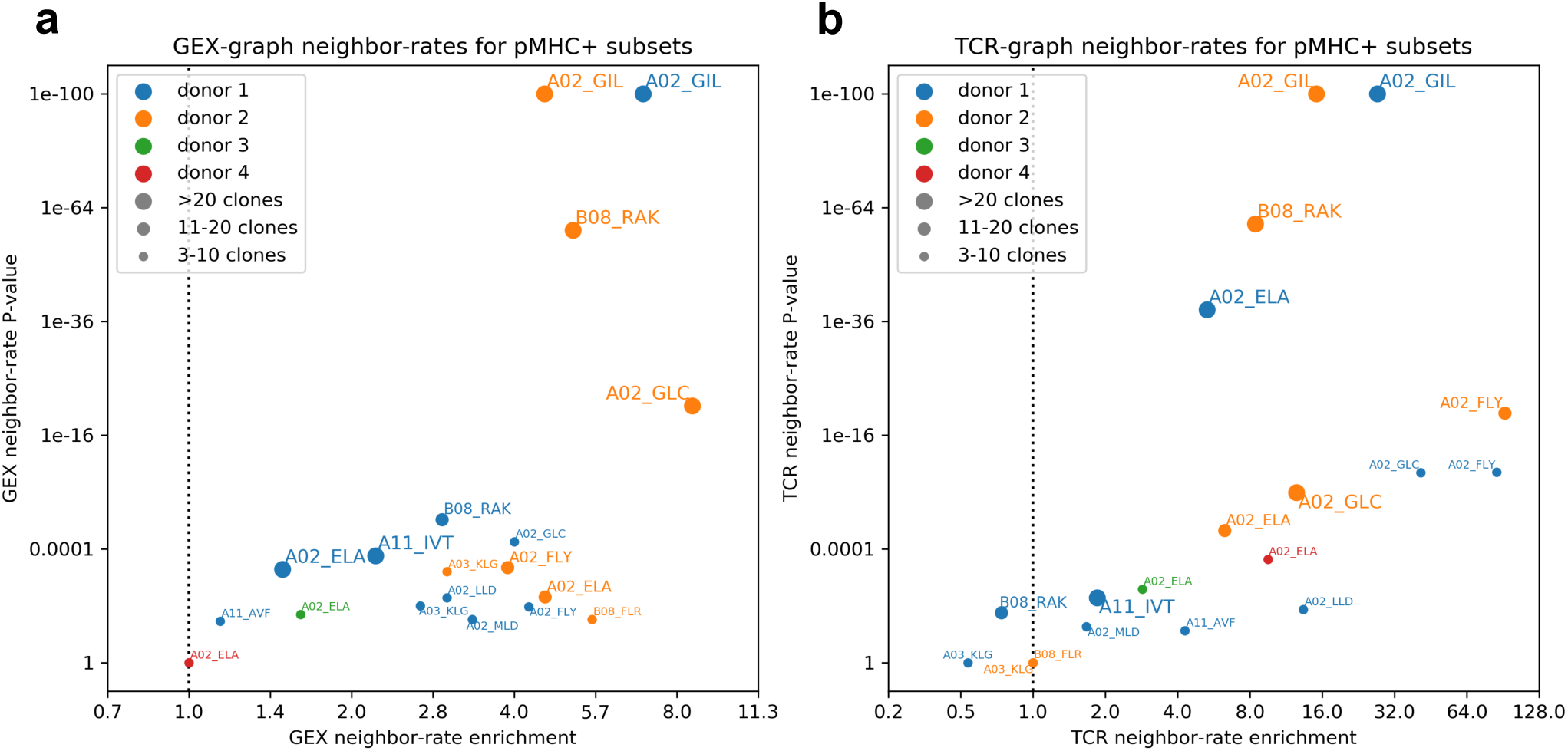
CoNGA identifies convergence of TCR sequence and gene expression profile within pMHC-positive clonotype subsets. Each marker represents a population of pMHC-positive clonotypes in one of the four *10x_200k* donors. Markers are labeled with the two-digit HLA allele and the first three amino acids of the peptide for the given pMHC (see **Table 4** for details); colors indicate the source donor and symbols are sized based on the number of pMHC+ clonotypes found as indicated in the legend. Markers are positioned based on the relative enrichment or depletion of GEX **(a)** or TCR **(b)** graph edges between pMHC-positive clonotypes compared to random expectation (*x*-axis; *>* 1 indicates enrichment while *<* 1 indicates depletion) and corresponding *P* value (*y*-axis).

We analyzed the expression patterns of specific marker genes to better understand these shared gene expression profiles. We performed all-against-all differential expression analyses to identify upregulated genes within each pMHC-positive subset. To visualize the results of this analysis, we selected the most common differentially expressed genes in these comparisons and created a gene expression heat map, clustering both the rows (pMHCs) and columns (genes) by similarity **(Fig. 10)**. Examination of the expression patterns in **Figure 10** reveals a number of trends: the naive responses (MART1 and B08_RAK in the B*08-negative donor 1) cluster together at the top and show higher levels of CD45RA and lower levels of *CCL5* and CD45RO; flu-M158 responses cluster together based on shared expression of specific markers including *GNLY, ITGB1*, and *IFITM1*; EBV-specific responses show what may be a partitioning based on whether the antigens are ‘early’ or ‘latent’ genes, with the early-gene responses showing higher *CCL5* and lower CD45RO compared to the ‘latent’-gene responses **(Fig. S11)**.

**Figure 10:**
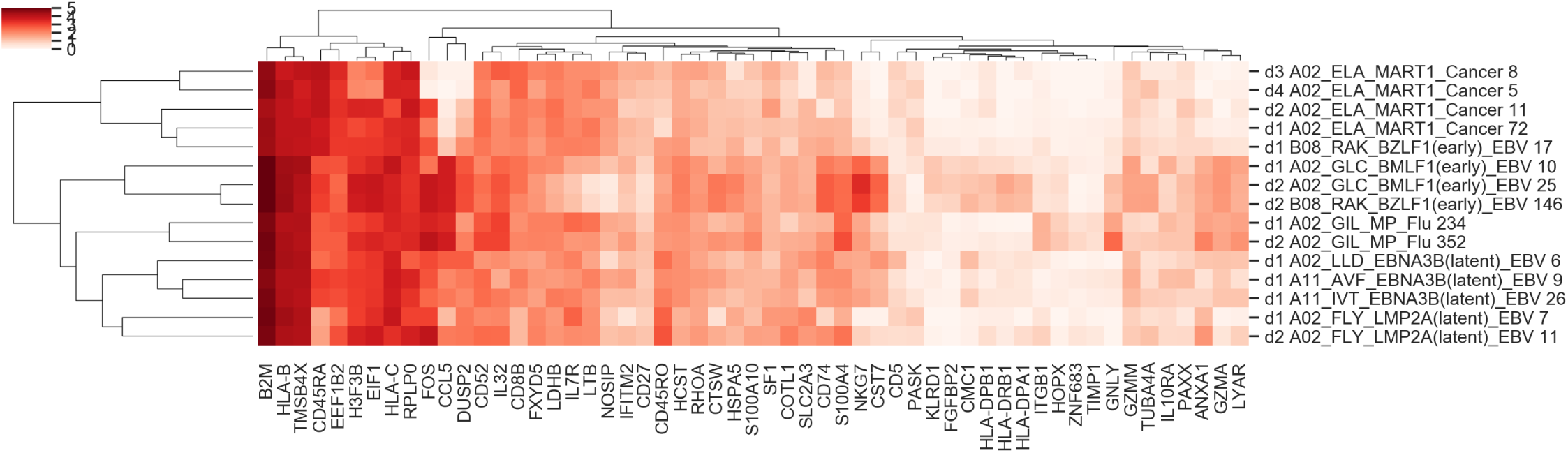
pMHC-positive clonotype populations show distinctive gene expression profiles. Each row corresponds to a pMHC-positive population; each row corresponds to a differentially expressed gene. Cells are colored according to the normalized transcript count (log+1 transformed) for the given gene in the corresponding population.

## Discussion

In this study, we have introduced and applied a new analytical tool, clonotype neighbor graph analysis or “CoNGA”, which we demonstrated to be capable of uncovering relationships within T cell populations defined by shared TCR sequence and gene expression features within large single-cell datasets. Previous works connecting the T cell state to its antigen-specificity have been limited to measuring variation in gene expression within cells of the same clonal lineage. CoNGA circumvents this strict limitation by defining neighborhoods based on TCR similarity then representing each clonal lineage within the neighborhood by a single representative cell, thus preserving phenotypic and TCR information from unexpanded clones that might otherwise be ignored. Application of CoNGA’s graph-vs-graph analysis on a diverse collection of datasets uncovered a number of previously unrecognized connections between TCR and GEX space, including distinct GEX profiles of epitope-specific T cells; bias in the repertoire selection of naive CD8+ and CD4+ T cell populations; multiple populations of thymic T cells with biased TCR repertoires; and a putative MHC-independent, *HOBIT/HELIOS*-expressing T cell subset detected both in the thymus and peripheral blood with distinctive CDR3 sequence features.

Further, while the identification of marker genes associated with cells clustered in GEX space is a routine part of single-cell analysis, there are currently no available methods for systematically identifying genes associated with TCR clusters or TCR sequence biases that define GEX clusters. CoNGA addresses this gap with its graph-vs-feature analysis by measuring a number of default TCR properties (amino acid composition, hydrophobicity, length, TCRdist score, etc.) before scanning the GEX space to detect clustered areas enriched for one or more of these features. Applying this mode of analysis revealed the long CDR3s of the *HOBIT*+ population enriched for hydrophobic residues, and a novel and highly significant correlation between expression of the *EPHB6* gene and usage of the *TRBV30* gene segment. Importantly, this mode of analysis is not limited to only TCR features but any other labelled feature (e.g. pMHC, cell surface marker, etc.) linked, quantified, and integrated into the dataset. In this regard, CoNGA analysis applied to a graph defined by single-cell pMHC-binding data determined that T cell populations specific for different pMHCs show distinctive GEX profiles, with evidence of clustering of EBV-epitope specific T cell populations according to the stage (early or latent) of the gene from which the epitope is derived. By systematically investigating connections between TCR sequence properties and GEX space within the dataset, CoNGA may significantly reduce the amount of time, effort, and missed correlations inherent in a manual approach. We believe that these findings and our analysis pipeline will be of interest to a range of researchers working on T cells and single-cell analysis.

An important next step will be to validate our findings by applying CoNGA to other datasets with GEX and TCR (and perhaps pMHC binding) information, as they become available. The rapid pace of single-cell technology development suggests that new and larger datasets, with additional phenotypic information from DNA-barcoded tagging reagents spanning diverse biological and clinical settings, will become available in the near future. It will also be important to experimentally characterize the T cell populations identified by CoNGA, which should be possible using flow cytometry and the marker genes highlighted by CoNGA clustering.

Our analysis has a number of important limitations that could be addressed in future work. First, a consequence of operating at the level of clonotypes rather than individual cells is that we miss out on variation within the cells of expanded clones. This includes variation in gene expression profile as well as variation in strength of pMHC binding (for datasets with pMHC multimer binding information). Although we found that gene expression was largely consistent within clonally related cells, it may be worth exploring approaches in which cellular resolution is preserved, for example by defining graphs at the level of individual cells and masking out intra-clonotype neighbor edges to eliminate the strong signal of clonal GEX/TCR correlation. It is also important to keep in mind that the results of applying CoNGA will depend critically on the distance measures used to define clonotype similarity and construct the neighbor graphs. Other measures of TCR similarity, for example derived from structural simulations, or of gene expression similarity, may highlight different features of GEX/TCR correlation and/or may be more sensitive. The same is true of other measures of TCR/GEX correlation, for example ones that directly use distances rather than neighbor graphs. In preliminary experiments we found neighbor-graph overlap to be generally superior to direct correlation of distance distributions **(Fig. S12)**, but there are many other possible approaches. Another limitation is that, in our experience, successful application of CoNGA requires a relatively large number of unique clones (at least several hundred), which depending on the degree of clonal expansion may require a substantially larger number of individual cells. Finally, the generality of the biological observations we report here should be weighed against the small number of donors examined. Future studies on larger cohorts will be necessary to definitively assess some of our observations.

To our knowledge, CoNGA is the first algorithm reported for the systematic detection of GEX/TCR correlation. As such, there are many possible extensions to explore in future work. CoNGA is agnostic to the source of the clonotype graphs, and hence could be applied to graphs defined by new similarity measures (based on surface protein expression, for example), new T cell clustering approaches ^31^, epigenetic rather than gene expression profiles, or new immunological and clinical phenotypes. CoNGA could also be applied to B cell clonotypes by incorporating a BCR sequence similarity score analogous to TCRdist. We applied CoNGA to identify correlation between numerical functions defined on TCR sequence (CDR3 amino acid properties and the iMHC score) or on the GEX profile (expression of individual genes), but other scalar functions of TCR/GEX/pMHC-binding could be used instead and might allow sensitive detection of specific axes of GEX/TCR correlation. Another exciting direction would be to use CoNGA to analyze the role of genetic variation outside the TCR region on GEX profile, TCR sequence, or pMHC binding: merging datasets from genetically diverse individuals and defining genotype similarity graphs might be one approach for doing this. Finally, it will be worthwhile to explore the use of more sophisticated graph-correlation algorithms developed in the computer science and machine learning communities as alternatives to the neighborhood-overlap and neighborhood-score enrichment that we have applied here.

The results of our analyses have a number of broader implications. First, the observation of a diversity of gene expression profiles across the different epitope-specific T cell populations argues for a broad continuum of memory T cell phenotypes ^32^ rather than a small number of discrete subsets. Indeed, the definition of memory phenotypes would seem to be significantly determined by the eliciting pathogen. It also suggests that improved prediction of target pMHC epitopes for T cells might be possible by combining TCR sequence with information on GEX profile ^33^. The putative MHC-independent and naive T cell populations identified by CoNGA hint at developmental influences of TCR sequence on T cell fate that go beyond the well-characterized role of invariant and semi-invariant TCRs ^34^. We are optimistic that new analytical approaches combined with novel high-throughput single-cell experiments will continue to illuminate new aspects of adaptive immunology in the coming years.

## Methods

### CoNGA software package

An open-source `python3` package implementing CoNGA graph-vs-graph and graph-vs-feature analysis is available from the software repository github (https://github.com/phbradley/conga). The `conga` package is built on the `scanpy` ^35^ python package (https://github.com/theislab/scanpy) for single-cell analysis and makes heavy use of `scanpy`’s `AnnData` object to store integrated gene expression and TCR sequence data. We are grateful to the authors of `scanpy` for creating such a robust and useful package. CoNGA includes an implementation of the TCRdist ^12^ distance calculation and TCR logo construction routines. Finally, CoNGA depends on the standard python data science tools `numpy, scipy, matplotlib, pandas,` and `scikit-learn` for visualization, data manipulation, and statistical calculations.

### TCR analysis

Clonotype data from 10x genomics is first converted into a TCRdist ‘clones file’ and the matrix of TCRdist distances is computed. By default, the 10x clonotype definitions are filtered to remove spurious chain sharing and merge split clonotypes (for example due to partial recovery of a second TCRalpha transcript). Kernel principal components analysis as implemented in `scikit-learn`’s `KernelPCA` class is then used to extract the top 50 components of variation from this distance matrix; these kernel PCs can be directly incorporated into the standard single-cell workflows for clustering and dimensionality reduction in place of the principal components extracted from the gene expression counts matrix. For generation of 2D landscape projections, CoNGA uses the UMAP algorithm for dimensionality reduction ^15^ as implemented in `scanpy.tl.umap`. Clusters of clonotypes with similar T cell receptor sequences are identified with the Louvain ^16,17^ graph-based clustering algorithm (`scanpy.tl.louvain`). Both UMAP projection and clustering rely on a nearest neighbors calculation conducted with the `scanpy.pp.neighbors` routine with 10 neighbors and 50 principal components (the 50 kernel PCs computed from the distance matrix). To annotate the Louvain clusters in CoNGA visualizations, the most frequent V segment in each cluster is identified and appended to the cluster name if it is present in at least 50% of the clustered TCRs, uppercased if present in at least 75% of the TCRs (clusters are initially named with consecutive integers, starting at 0 with the largest cluster).

### TCR sequence features

For each clonotype, CoNGA calculates a set of TCR sequence-based scores for use in graph-vs-feature analysis and for annotating graph-vs-graph cluster pairs (**Table S1**). First, a set of 28 different amino acid properties **(Fig. S8, Table S1)** are averaged over the central amino acids in the alpha and beta chain CDR3 loops (excluding the first 4 and last 4 residues of each CDR3, where the full CDR3 sequence is defined as beginning with the conserved cysteine and ending with, and inclusive of, the phenylalanine immediately before the GXG motif in the J region). These scores include a set compiled from original sources ^36–41^ by the authors of the VDJtools package ^42^ as well as the five Atchley factors ^43^. Seven additional sequence-based scores are calculated: ‘alphadist’, which measures the ordinal distance between the Valpha and Jalpha genes when the full set of gene segments is ordered by genomic position; ‘imhc’, the iMHC score (detailed below); ‘cd8’, a simple CD8-versus-CD4 preference score calculated from the TCR V and J gene usage, CDR3 length, and CDR3 amino acid composition, based on frequency differences between flow-sorted CD8+ and CD4+ TCR sequence repertoires (AS, unpublished results); ‘cdr3len’, total CDR3 length; ‘mait’, which assigns a score of 1 to TCRs with an alpha chain using the TRAV1-2 and TRAJ33/TRAJ20/TRAJ12 segments (TRAV1 and TRAJ33 in mouse) and a CDR3 length of 12, and 0 to all other TCRs; ‘inkt’, which assigns a score of 1 to TCRs with the TRAV10/TRAJ18/TRBV25 gene combination and a CDR3 length of 14, 15, or 16 (TRAV11/TRAJ18 and length 15 for mouse); and ‘nndists_tcr’, which measures the density of TCR sequences nearby the scored clonotype by calculating the average TCR distance to the nearest 1% of clonotypes. The iMHC (for ‘independent of pMHC’) score is a weighted linear combination of TCR sequence features (**Table S2**). The parameters were fit by using L1-regularized logistic regression to discriminate the TCR sequences of *HOBIT*+ CoNGA hits (CoNGA score<0.2) in GEX cluster 2 of dataset *10x_200k_donor1* **(Fig. 3)** from the TCRs of the clonotypes in the other GEX clusters. We chose to draw the background clonotypes exclusively from the other GEX clusters to avoid inclusion of genuine *HOBIT*+ TCR sequences in our negative set.

### Gene expression analysis

Gene expression data in the form of read count matrices are processed according to standard workflows implemented in `scanpy` to eliminate cells and genes with low counts, high mitochondrial content, etc. Variable genes are identified and principal components analysis (PCA) is used to project the high-dimensional gene expression data down to a smaller set of components per cell (the default is 40 components). These gene expression PCs are used to select a single representative cell for each clonotype by taking the cell with the smallest average Euclidean distance in PC space to the other cells in the clonotype. Once the dataset has been reduced to a single cell per clone, the UMAP and Louvain clustering tools are applied to the PCA matrix to produce a gene expression landscape and a set of gene expression clonotype clusters. DEGs in clonotype groupings (for example the set of CoNGA hits in a cluster pair) are identified using the `sc.tl.rank_genes_groups` routine with the ‘wilcoxon’ method.

The large thymus atlas T cell dataset ^27^ combined a heterogeneous set of donors and samples; merging these data to generate integrated projections and clusters required the original authors to perform an iterative batch correction scheme. As it was not immediately obvious how to recover the processed gene expression components from the publicly available data, and as a test of CoNGA’s robustness to alternative neighbor graphs, we elected to use the provided 3D UMAP coordinates in lieu of gene expression PCs for the CoNGA GEX neighbor calculations described below. We also directly borrowed the GEX clusters from the original paper rather than reclustering the dataset.

### Graph-vs-graph correlation analysis

In CoNGA graph-vs-graph correlation analysis, similarity graphs defined by gene expression and by TCR sequence are compared to identify vertices (clonotypes) whose neighbor sets in the two graphs overlap significantly. The CoNGA score assigned to a clonotype equals the probability of seeing an equal or larger overlap between its GEX and TCR neighborhoods by chance, multiplied by the total number of clonotypes to correct for multiple testing. The hypergeometric distribution is used to estimate this probability, as implemented in the `scipy.stats` module. Two types of similarity graphs can be used in CoNGA: K nearest neighbor (KNN) graphs, in which each clonotype is connected to its K nearest neighbors in gene expression or TCR space (**Fig. 1a**); and *cluster graphs*, in which each clonotype is connected to all the clonotypes in the same (GEX or TCR) cluster. The neighbor number K for constructing KNN graphs is specified as a fraction of the total number of clones; for the calculations reported here, neighbor fractions of 0.01 and 0.1 were used. The CoNGA score assigned to a clonotype is the minimum score over all graph comparisons, of which there were 6 combinations in the calculations reported here (GEX_KNN vs TCR_KNN, GEX_KNN vs TCR_cluster, and GEX_cluster vs TCR_KNN, for both the 0.01 and 0.1 KNN neighbor fractions). Although in principle taking a minimum could lead to inflation of the significance scores, we find in practice from shuffling experiments that a threshold of 1.0 on the CoNGA score remains a useful indicator of genuine signal, particularly when we focus on cluster pairs with a minimum number of CoNGA hits. This may reflect correlation between neighborhoods of nearby clonotypes, which reduces the effective multiple-testing burden.

### Graph-vs-feature correlation analysis

In CoNGA graph-vs-feature correlation analysis, numerical features defined on the basis of one property (GEX or TCR) are mapped onto similarity graphs defined by the other property, and graph neighborhoods with biased score distributions are identified. As GEX properties we consider the expression levels of all the individual genes as well as a feature (‘nndists_gex’) that captures the density of nearby clonotypes by calculating the average distance in GEX space to the nearest 1% of the clonotypes. The TCR features were described in an earlier section. As this analysis involves a large number of differential expression calculations (roughly the number of clonotypes times the number of different similarity graphs times the number of features), we use a two-step procedure that combines a pre-filter with the t-test followed by the more time-intensive Mann-Whitney-Wilcoxon (MWW) calculation for the top 100 hits per clonotype and graph that pass a t-test significance threshold ten times higher than the target threshold. The final significance score assigned to a detected association equals the raw MWW P-value multiplied by the product of the number of clonotypes and the number of features, to correct for multiple testing.

### Analysis of pMHC binding

In the *10x_200k* experiment, T cells were stained with a panel of 50 DNA-barcoded pMHC multimer reagents. Sequence reads for each of the pMHC barcodes were counted along with the reads for intracellular transcripts and included in the raw count matrix provided by 10x Genomics. The first step in our analysis was to assign individual T cells and T cell clonotypes as positive for binding to specific pMHC multimers based on the observed read counts for the pMHC DNA barcodes. A cell was called positive for the pMHC multimer with the highest barcode count if the natural logarithm of that pMHC’s barcode count exceeded the next highest log-count by at least 2.0 (corresponding to a fold-difference in barcode counts of roughly 7.5; all counts were augmented by 1 prior to taking logarithms). To assign clonotypes to pMHCs, we averaged the log-counts for each pMHC over all the cells in the clonotype and again applied a threshold of 2.0 between the top and second-highest averaged-log-counts. The results of this pMHC-binding analysis are summarized in **Table 4** for all pMHCs with at least 5 positive clones in one of the four samples. We can see that, with the exception of the ‘sticky’ pMHC A03_KLG, the majority of pMHC+ positive cells belong to clones that are also called positive.

### Flow Cytometry Analysis and qRT-PCR of *HOBIT*+ population

To identify the *HOBIT*+, iMHC population of CD8 T cells using cell surface markers we performed flow cytometric analysis on PBMCs collected from apheresis rings of blood donors. PBMC samples were blocked with TruStain FcX prior to staining for CD3 (APC-Fire750, SK7), CD4 (PE-Cy7, OKT4), CD56 (PE, 5.1H11), CCR7 (BrilliantViolet 785, G043H7), CD11b (BrilliantViolet 711, ICRF44), CD45RA (BrilliantViolet 421, HI100), CD45RO (BrilliantViolet 605, UCHL1) (BioLegend), CD8A (violetFluor450, RPA-T8), CD14 (biotin, 61D3), CD19 (biotin, SJ25C1), CD16 (biotin, 3G8) (Tonbo Bioscience), KIR2D (FITC, NKVFS1), KLRC2 (PE-Vio615, REA205) (Miltenyi), CD8B (PerCP-eFluor710, SIDI8BEE, Invitrogen), and CD248 (AlexaFluor 647, B1/35, BD Biosciences) for 30’ at RT in PBS containing 2% FCS and 1 mM EDTA prior to secondary staining with streptavidin-BrilliantViolet 510 (Biolegend) for 15’ on ice. Stained cells were then analyzed with an Aurora spectral analyzer (Cytek) or sorted by an iCyt (Sony). Analysis of flow cytometry data was performed with FlowJo (BD Biosciences).

To confirm expression of genes associated with the *HOBIT*+ CoNGA population, for four donors the PBMCs were sorted into three populations: KLRC2+ KIR2+/- (dump-, CD3+, CD56+/-, CD8+, CD45RA+, CD45ROdim/-, CD248-, CCR7-, KLRC2+, KIR2D +/-), KLRC2-KIR2+ (dump-, CD3+, CD56+/-, CD8+, CD45RA+, CD45ROdim, CD248-, CCR7-, KLRC2-, KIR2D+), and CD45RO+ (dump-, CD3+, CD56+/-, CD8+, CD45RA-, CD45RO+) as a control for assessing enrichment of the signature genes. Total RNA was extracted from the sorted cells with RNeasy Micro Columns (Qiagen), converted into cDNA (iScript, Bio-Rad), and assayed for ZNF683, KLRC2, KLRC3, and GAPDH expression using gene-specific primers and SYBR Green chemistry (iTaq, Bio-Rad) by qRT-PCR on a CFX96 (Bio-Rad). The fold-change relative to the CD45RO+ population was calculated using ΔΔCt with GAPDH as the housekeeping gene. The following are the primer sequences: KLRC2_Fwd:CCTGATGGCCACTGTGTTAAA,KLRC2_Rev:GCGTTCTTGTATTCGGGGAA, KLRC3_Fwd:CAGGCCTGTGCTTCAAAGAA, KLRC3_Rev:GAAACACACCAATCCATGAGGAA, ZNF683_Fwd:CAAAGCGGGTCCCATTGAGTT, ZNF683_Rev:TGCACTCGTACAGGATTTTGC.

## Acknowledgments

The authors would like to thank Jongeun Park and Sarah Teichmann for assistance with the thymus atlas T cell dataset, Erick Matsen for comments and suggestions on an earlier version of this manuscript, Evan Newell and Timothy Bi for helpful discussions, and Nicholas Bradley for suggesting the use of kernel principal components analysis. We would also like to thank the developers of the `scanpy` single-cell analysis package, which provides the framework on which the CoNGA software is built. This research was supported by NIH grant R01 AI136514 to PT, the St. Jude Neoma Boadway Postdoctoral Fellowship to SS, and ALSAC (PT).

## Figure Legends

**Supplementary Figure 1. T cells belonging to the same clonotype have similar gene expression profiles.** Gene expression UMAP projections of the *10x_200k_donor2a* dataset before condensing to a single cell per clonotype, with the 16 largest clonotypes shown in blue (one per panel) and the remainder of the dataset in gray.

**Supplementary Figure 2. CoNGA graph-vs-graph analysis of human and mouse peripheral blood T cells.** CoNGA graph-vs-graph results for two additional PBMC T cell datasets: **(a-c)** human CD4 and CD8 T cells (*vdj_v1_hs_pbmc3*); **(d-f)** mouse CD4 and CD8 T cells (*vdj_v1_mm_balbc_pbmc*). Same arrangement of plots as in main text **Figure 2**.

**Supplementary Figure 3. Specific versus non-specific binding in the *10x_200k* dataset.** Comparison of binding data for four ‘specific’ pMHC multimers (A02_GIL, A02_ELA, B08_RAK, A02_GLC) and four ‘sticky’ pMHC multimers (A03_KLG, A03_RLR, A03_RIA, A11_AVF) in the *10x_200k_donor2* dataset. **(a)** GEX landscapes colored by pMHC binding signal (log(barcode_read_count+1)). **(b)** TCR landscapes colored by pMHC binding signal. The ‘specific’ pMHCs show binding that is focused in specific areas of the landscapes, whereas the binding of the putative ‘sticky’ pMHCs is dispersed across the landscapes. **(c)** The Pearson correlation between binding profiles for different pMHCs is shown in matrix form according to the indicated color mapping. The specific pMHCs show very little correlation whereas the sticky pMHCs are significantly correlated in their binding, suggesting that a shared cellular property (TCR or CD8 surface expression, general level of activation) is jointly influencing their binding. Note that A11_AVF (and A11_IVT) show additional specific binding in donor 1, who is A*11:01 positive; the A*03:01 pMHC multimers appear non-specific regardless of donor HLA type (data not shown).

**Supplementary Figures 4-7.** CoNGA graph-vs-graph results for *10x_200k_donor1 - 10x_200k_donor4*, showing logos for all cluster pairs of size>=5. For each cluster pair, the top two pMHCs and their average (log) read counts are shown in the cluster pair dendrogram below the cluster pair sizes. Note that the top pMHCs for the *HOBIT*+ cluster pairs in donor 1 belong to the set of sticky pMHCs shown in **Figure S3**.

**Supplementary Figure 8. The 28 amino acid property scores currently used in CoNGA analyses. (a)** Clustering dendrogram (left) and matrix visualization of the amino acid property scores, normalized to have mean 0 and variance 1. **(b)** The correlation matrix used to construct the dendrogram in (a). Each entry is colored by the absolute value of the Pearson correlation coefficient of the property values for the corresponding row and column.

**Supplementary Figure 9. Gating strategy for KLRC2+ KIR2D+/- and KLRC2-KIR2D- CD8 T cells in Figure 5.**

**Supplementary Figure 10. Single-chain iMHC score distributions for TCR subsets.** Score distributions for CDR3α repertoires are shown on the left and for CDR3β repertoires on the right. Single-chain variants of the iMHC score were fit with L1-regularized logistic regression just as for the paired iMHC score. Subset labels are as follows: ‘CD1b’, GMM:CD1b-tetramer sorted T cells from Ref ^44^; ‘VDJdb-MHC1’, TCRs reported to bind to MHC class 1 presented epitopes in the VDJdb database ^45^; ‘VDJdb-MHC2’, TCRs reported to bind to MHC class 2 presented epitopes in the VDJdb database; ‘dMAIT’, diverse MAIT TCR sequences from Ref ^46^; ‘hobit’, T cells belonging to the *HOBIT*+/HELIOS+ population in *10x_200k* donors 1, 3, and 4.

**Supplementary Figure 11. Epitope-specific T cell populations differ in CD45RA/RO expression levels.** Log-transformed read counts for DNA-barcoded anti-CD45RA (x-axis) and anti-CD45RO (y-axis) antibodies, averaged over pMHC+ clonotypes, are plotted for the pMHCs shown in **Figure 10**. In the panel on the left, clonotypes are weighted equally, while in the panel on the right, larger clonotypes are given more weight (proportional to the logarithm of the clone size) to better reflect the underlying distribution of cells (particularly for the d1_A11 pMHCs, both of which have a relatively large number of positive cells distributed unevenly among a small number of clonotypes).

**Supplementary Figure 12. Comparison of graph-based and distance-based measures for assessing GEX/TCR correlation.** Comparison of CoNGA scores to a distance-based score (‘distcorr’) that measures, for each clonotype, the degree of correlation between the GEX and TCR distances from that clonotype to all other clonotypes in the dataset. Correlation is assessed using the Pearson correlation coefficient and associated P-value as returned by the `scipy.stats.linregress` function. **(a)** Scatter plots directly comparing the (negative log10-transformed) significance scores assigned to each clonotype in the datasets. The majority of points with significant *P* values lie below the *y=x* line, indicating that the CoNGA graph-overlap measure assigns a higher significance score than distance correlation. One difficulty with raw distance correlation is that it doesn’t discriminate between a set of clonotypes with low distances for both measures (nearby in GEX and in TCR space), on the one hand, and a set of clonotypes with high distances for both measures (far away in GEX and in TCR space), on the other: both increase the correlation coefficient, so a tight cluster in GEX and TCR space (like MAIT cells) can artificially elevate distcorr scores for distant clones. (b) CoNGA and distcorr scores mapped to the GEX and TCR landscapes for *10x_200k_donor2a*. The red ellipses indicate the A*02:M1_58_ clones, which appear to be completely missed by the distcorr measure.

---

**Table S1:** TCR sequence feature descriptions

| Feature<sup>a</sup> | Description |
| --- | --- |
| alphadist | Ordinal distance between the Vα and Jα gene segments when the TCRα locus is ordered by genomic position |
| cd8 | CD8 vs CD4 preference score |
| cdr3len | Combined length of the CDR3α and CDR3β regions |
| inkt | 1 for TCRs matching an iNKT sequence consensus, 0 otherwise |
| mait | 1 for TCRs matching a MAIT sequence consensus, 0 otherwise |
| imhc | MHC-independent score fit to discriminate the Hobit+/Helios+ TCRs |
| nndists_tcr | TCR neighborhood distance score equal to average TCRdist to nearest 1% of clonotypes in dataset |
| alpha<sup>36</sup> | Preference to appear in alpha helices |
| beta<sup>36</sup> | Preference to appear in beta sheets |
| turn<sup>36</sup> | Preference to appear in turns |
| surface<sup>39</sup> | Frequency in protein surface away from protein-protein interfaces |
| rim<sup>39</sup> | Frequency in rim region of protein-protein interfaces |
| core<sup>39</sup> | Frequency in core region of protein-protein interfaces |
| disorder<sup>40</sup> | Disorder-promoting amino acid score |
| charge | Amino acid charge |
| pH | Amino acid pH level |
| polarity | Polar/non-polar amino acids |
| hydropathy | Amino acid hydropathy |
| volume | Amino acid volume |
| strength<sup>38</sup> | Strongly-interacting amino acids as defined by analysis of MJ statistical potential |
| mjenergy<sup>37</sup> | Mean value of MJ statistical potential for each amino acid |
| kf1...kf10<sup>41</sup> | Values of 10 Kidera factors summarizing physicochemical properties of amino acids |
| af1...af5<sup>43</sup> | Values of 5 Atchley factors summarizing physicochemical properties of amino acids |

<sup>a</sup> Features 'alpha' through 'kf10' were compiled by the authors of the VDJtools<sup>42</sup> package

**Table S2:** iMHC score coefficients

| Feature | Coefficient | Std. coefficient<sup>a</sup> | Std. dev.<sup>b</sup> |
| --- | --- | --- | --- |
| len_AB | 0.368 495 | 1.013 617 | 2.750 693 |
| len_B | 0.321 411 | 0.606 267 | 1.886 267 |
| charge_AB | 0.603 610 | 0.137 170 | 0.227 249 |
| charge_B | 0.902 453 | 0.136 966 | 0.151 771 |
| volume_AB | 0.006 965 | 0.129 819 | 18.638 504 |
| Wfrac_AB | 4.141 282 | 0.117 301 | 0.028 325 |
| disorder_AB | −0.290 458 | −0.114 851 | 0.395 414 |
| volume_B | 0.009 481 | 0.114 739 | 12.102 429 |
| arofrac_AB | 1.217 256 | 0.093 160 | 0.076 533 |
| surface_B | −15.068 416 | −0.088 320 | 0.005 861 |
| Cfrac_B | 5.998 254 | 0.032 482 | 0.005 415 |
| Kfrac_AB | 0.441 505 | 0.011 270 | 0.025 526 |
| Cfrac_AB | 2.313 601 | 0.009 199 | 0.003 976 |
| Ffrac_AB | 0.013 840 | 0.000 550 | 0.039 716 |
| model_intercept | −41.230 464 |  |  |

<sup>a</sup> Coefficient scaled by feature standard deviation<sup>b</sup> Feature standard deviation over the *10x\_200k\_donor1* dataset.

**Figure S1:**
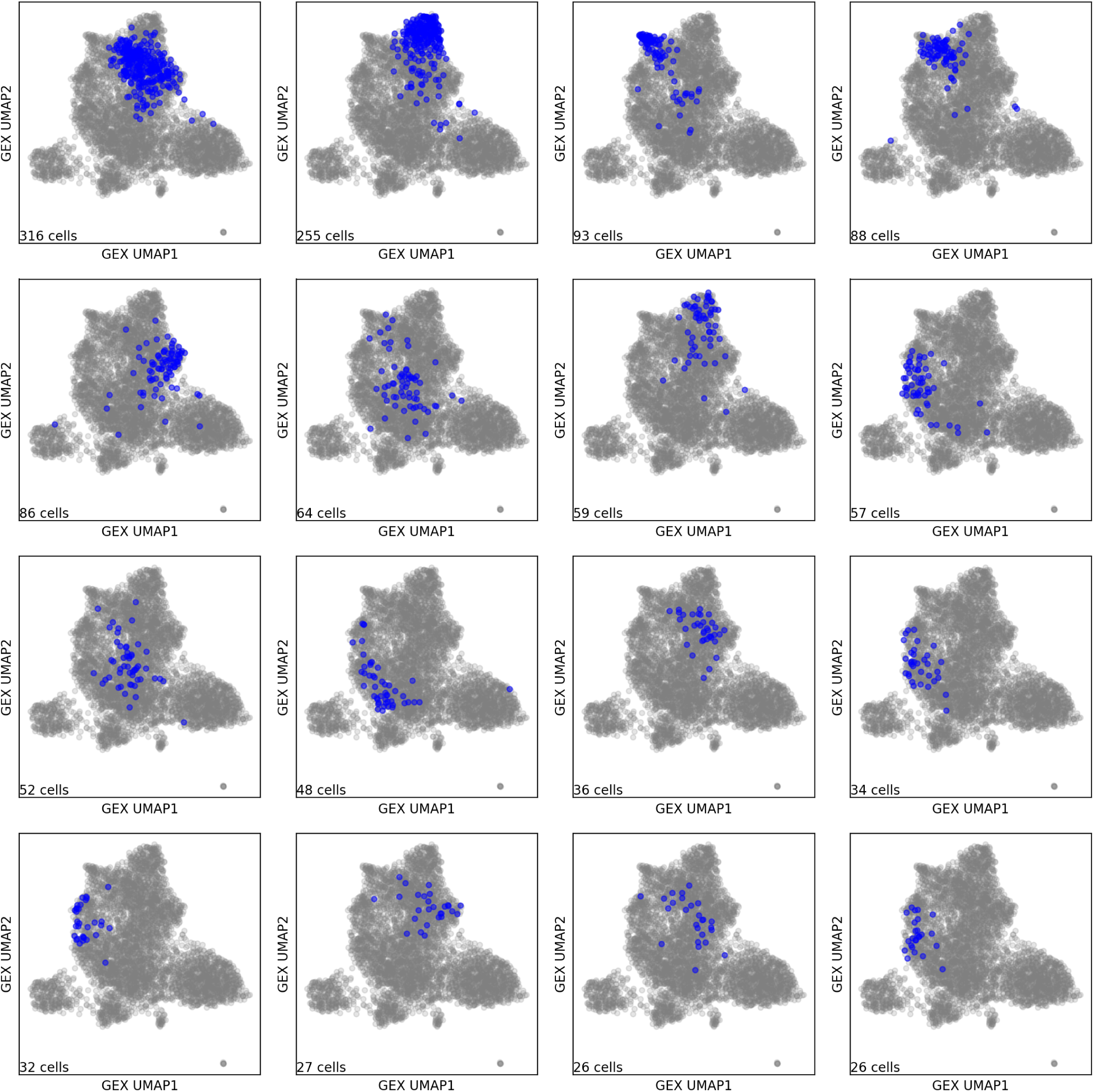
T cells belonging to the same clonotype have similar gene expression profiles. Gene expression UMAP projections of the *10x_200k_donor2a* dataset before condensing to a single cell per clonotype, with the 16 largest clonotypes shown in blue (one per panel) and the remainder of the dataset in gray.

**Figure S2:**
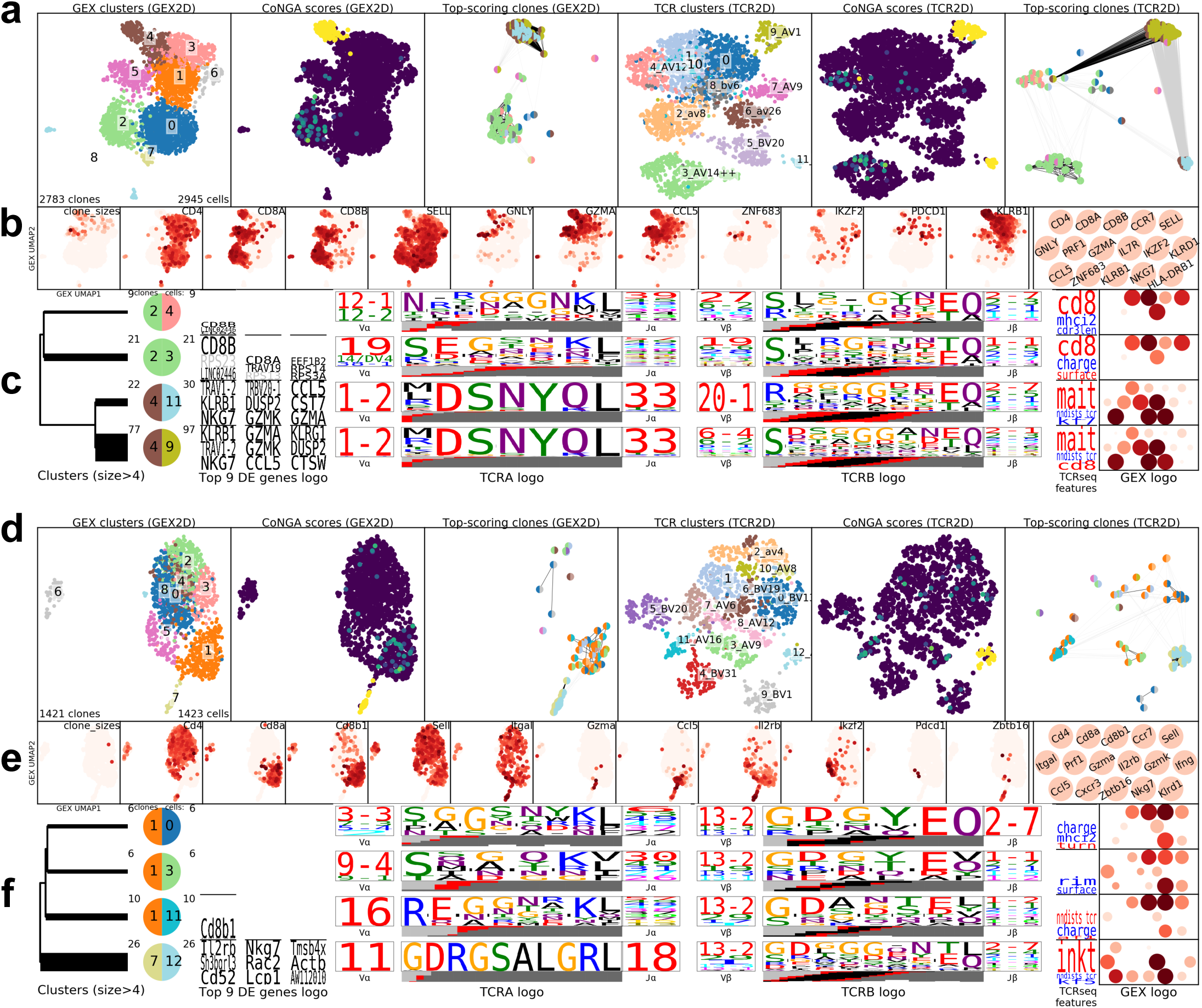
CoNGA graph-vs-graph analysis of human and mouse peripheral blood T cells. CoNGA graph-vs-graph results for two additional PBMC T cell datasets: **(a-c)** human CD4 and CD8 T cells (*vdj_v1_hs_pbmc3*); **(d-f)** mouse CD4 and CD8 T cells (*vdj_v1_mm_balbc_pbmc3*). Same arrangement of plots as in main text **Figure 2**.

**Figure S3:**
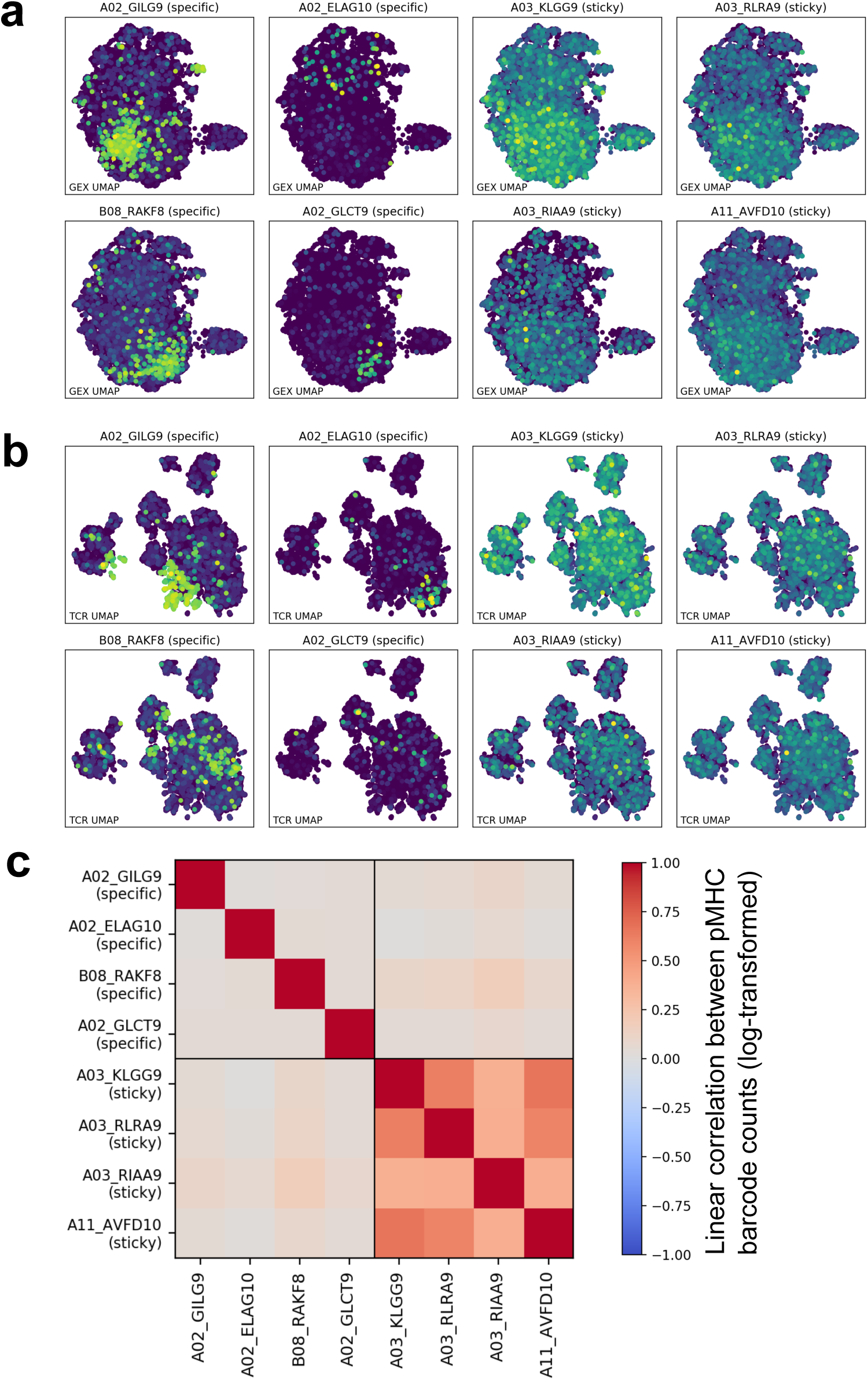
Specific versus non-specific binding in the *10x_200k* dataset. Comparison of binding data for four ‘specific’ pMHC multimers (A02_GIL, A02_ELA, B08_RAK, A02_GLC) and four ‘sticky’ pMHC multimers (A03_KLG, A03_RLR, A03_RIA, A11_AVF) in the *10x_200k_donor2* dataset. **(a)** GEX landscapes colored by pMHC binding signal (log-transformed barcode read counts). **(b)** TCR landscapes colored by pMHC binding signal. The ‘specific’ pMHCs show binding that is focused in specific areas of the landscapes, whereas the binding of the putative ‘sticky’ pMHCs is dispersed across the landscapes. **(c)** The Pearson correlation between binding profiles for different pMHCs is shown in matrix form according to the indicated color mapping. The specific pMHCs show very little correlation whereas the sticky pMHCs are significantly correlated in their binding, suggesting that a shared cellular property (TCR or CD8 surface expression, general level of activation) is jointly influencing their binding. Note that A11_AVF (and A11_IVT) show additional specific binding in donor 1, who is A*11:01 positive; the A*03:01 pMHC multimers appear non-specific regardless of donor HLA type (data not shown).

**Figure S4:**
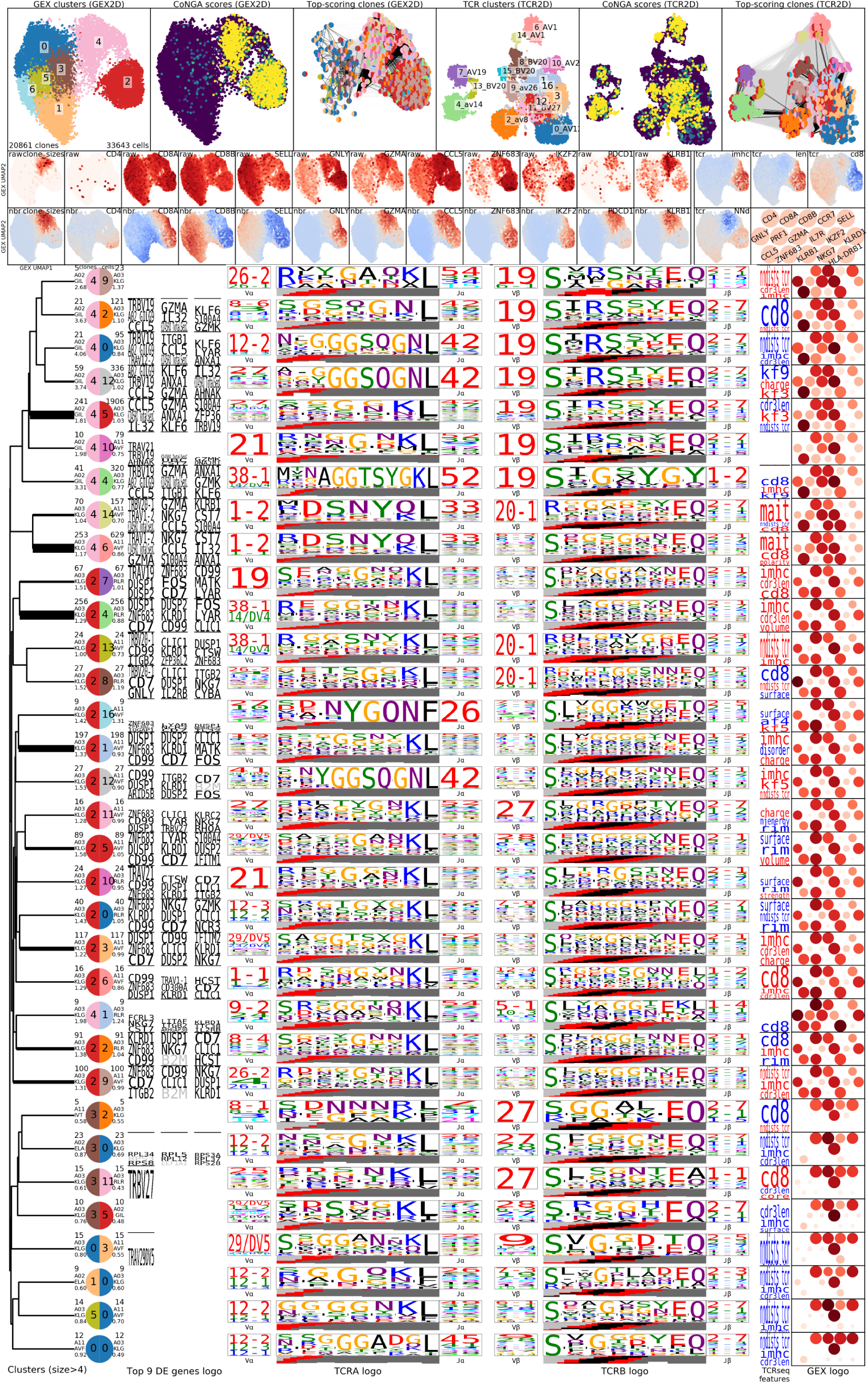
CoNGA graph-vs-graph results for the *10x_200k_donor1* dataset

**Figure S5:**
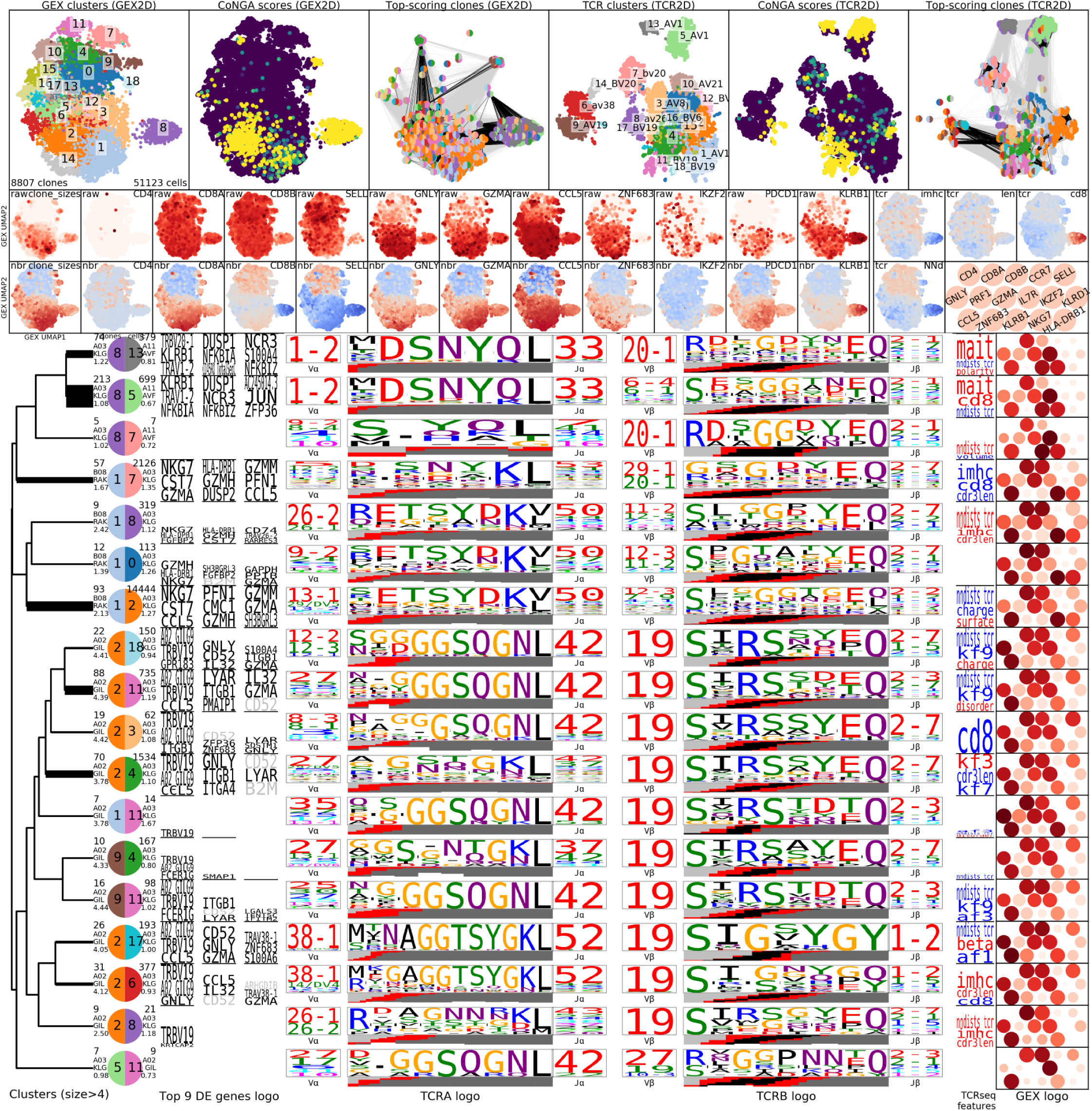
CoNGA graph-vs-graph results for *10x_200k_donor2* dataset. Logos are shown for all cluster pairs with at least 5 CoNGA hits. For each cluster pair, the top two pMHCs and their average (log) read counts are shown in the cluster pair dendrogram below the cluster pair sizes.

**Figure S6:**
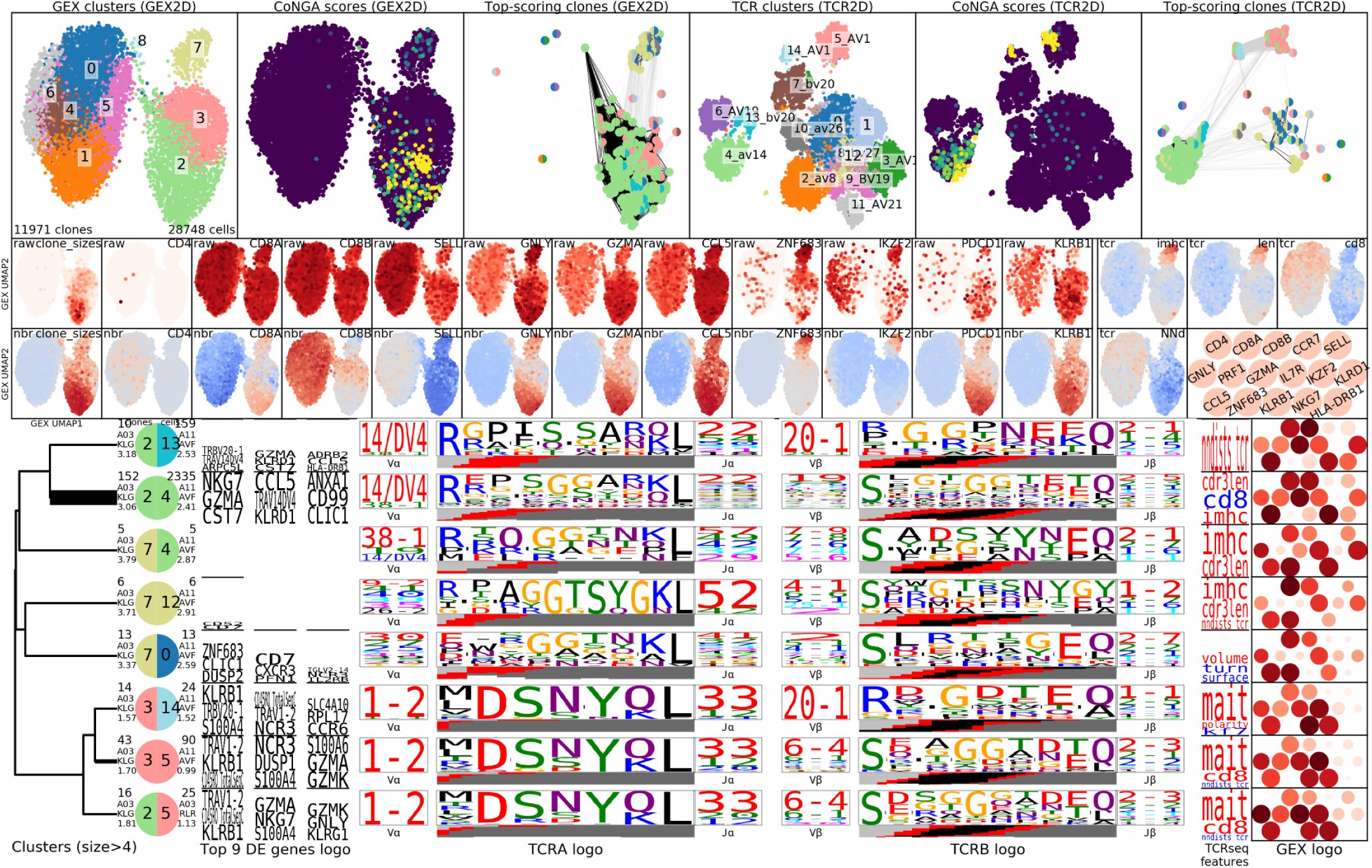
CoNGA graph-vs-graph results for *10x_200k_donor3* dataset. Logos are shown for all cluster pairs with at least 5 CoNGA hits. For each cluster pair, the top two pMHCs and their average (log) read counts are shown in the cluster pair dendrogram below the cluster pair sizes.

**Figure S7:**
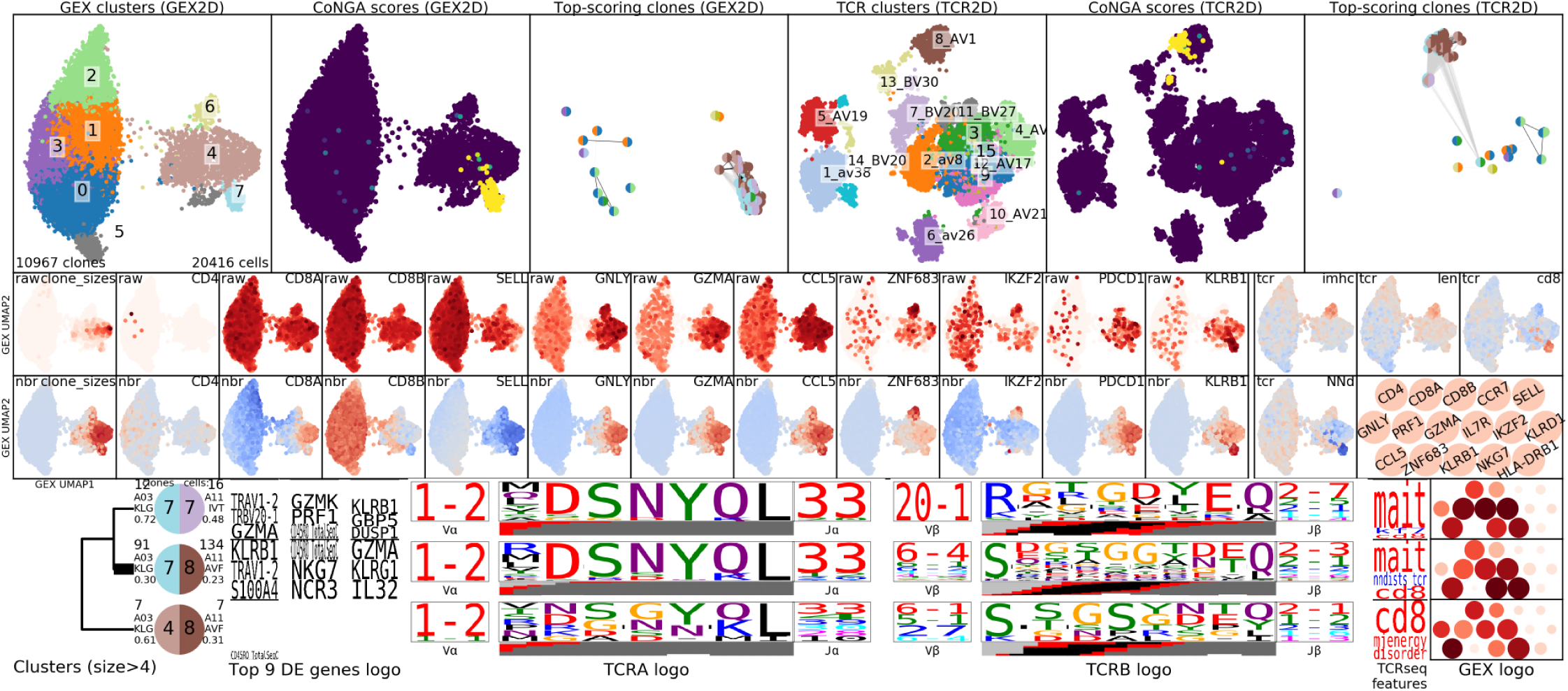
CoNGA graph-vs-graph results for *10x_200k_donor4* dataset. Logos are shown for all cluster pairs with at least 5 CoNGA hits. For each cluster pair, the top two pMHCs and their average (log) read counts are shown in the cluster pair dendrogram below the cluster pair sizes.

**Figure S8:**
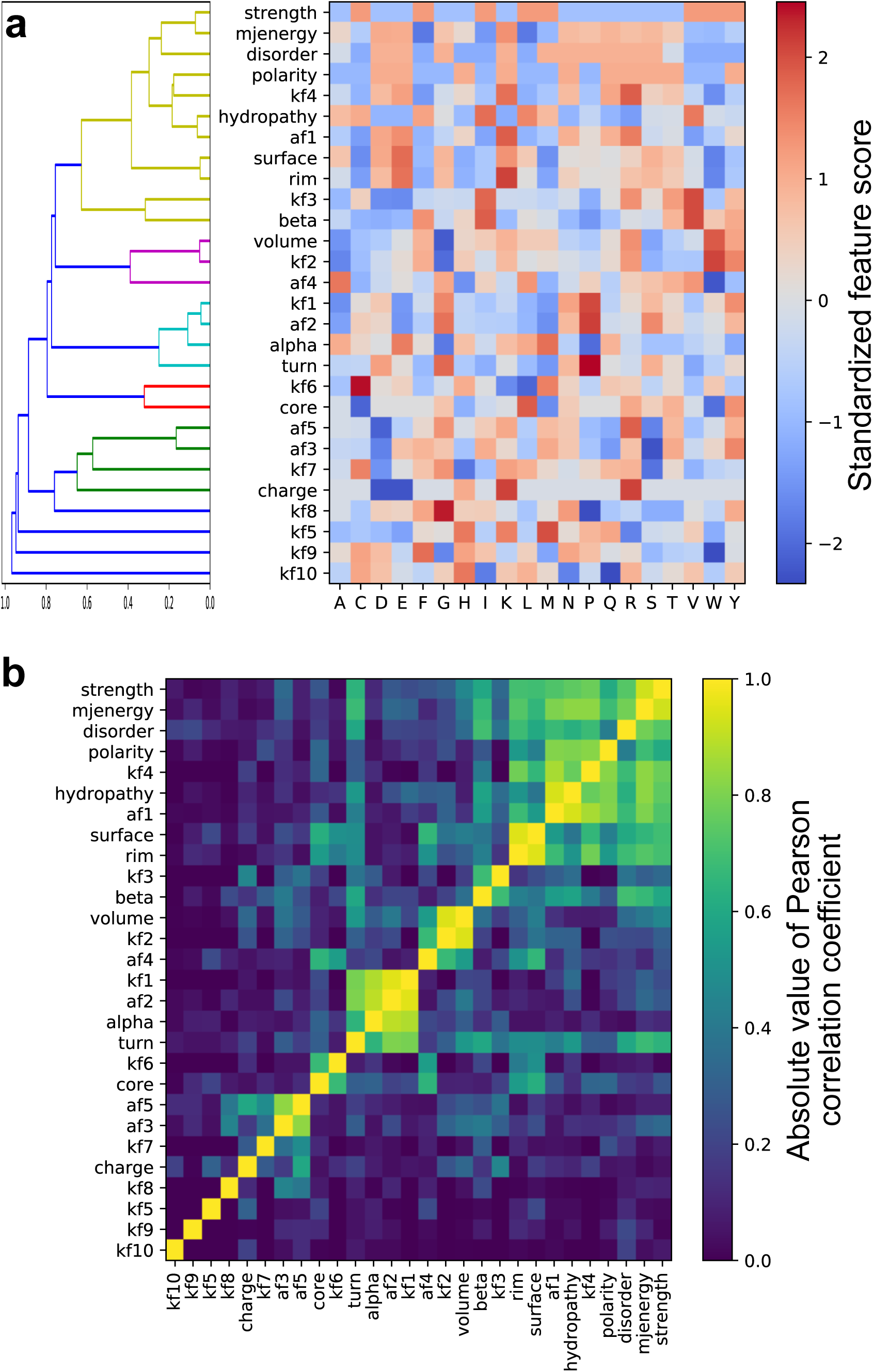
The 28 amino acid property scores currently used in CoNGA analyses. **(a)** Clustering dendrogram (left) and matrix visualization of the amino acid property scores, normalized to have mean 0 and variance 1. **(b)** The correlation matrix used to construct the dendrogram in (a). Each entry is colored by the absolute value of the Pearson correlation coefficient of the property values for the corresponding row and column.

**Figure S9:**
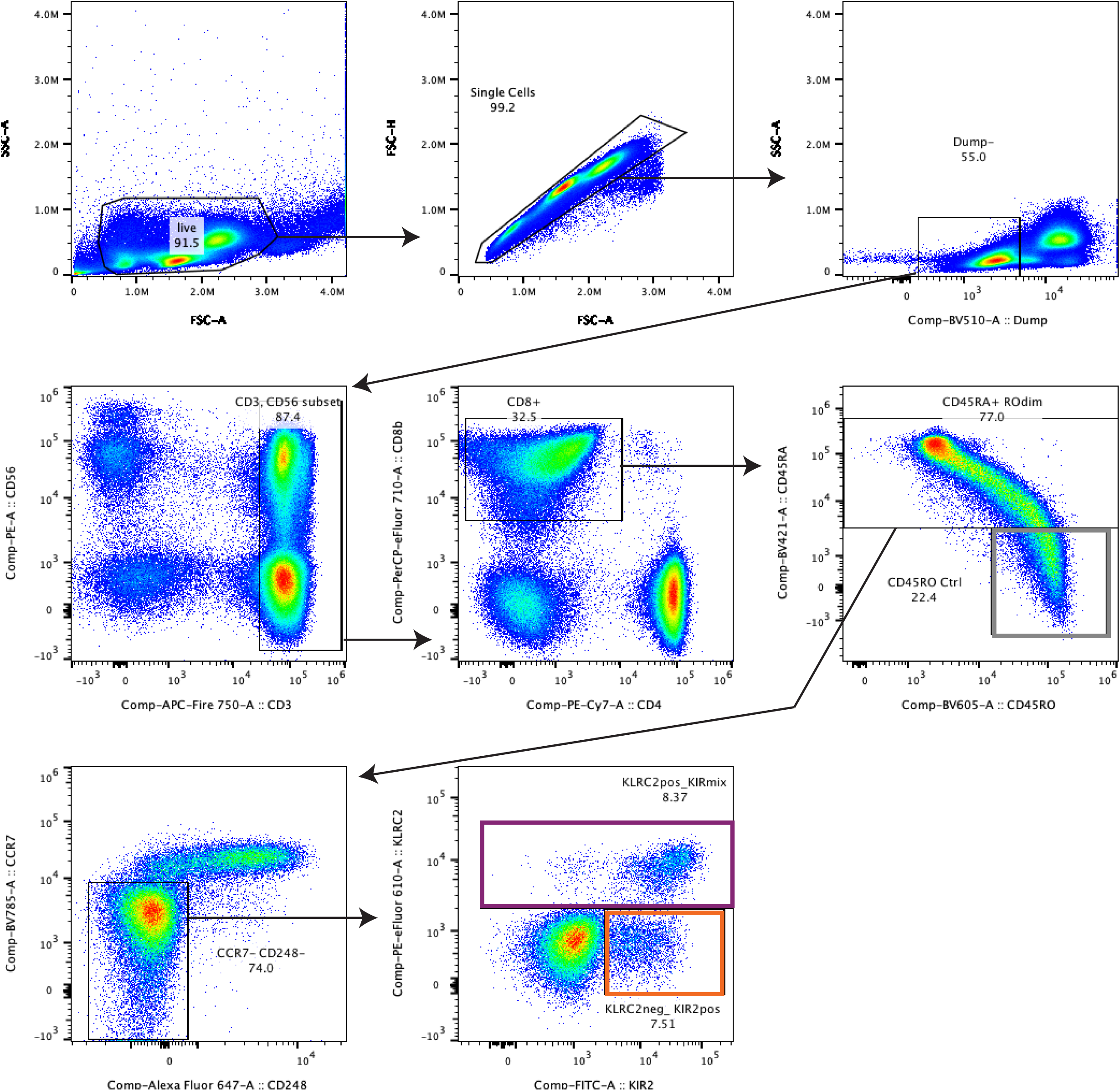
Gating strategy for KLRC2+ KIR2D+/- and KLRC2- KIR2D- CD8 T cells in Figure 5.

**Figure S10:**
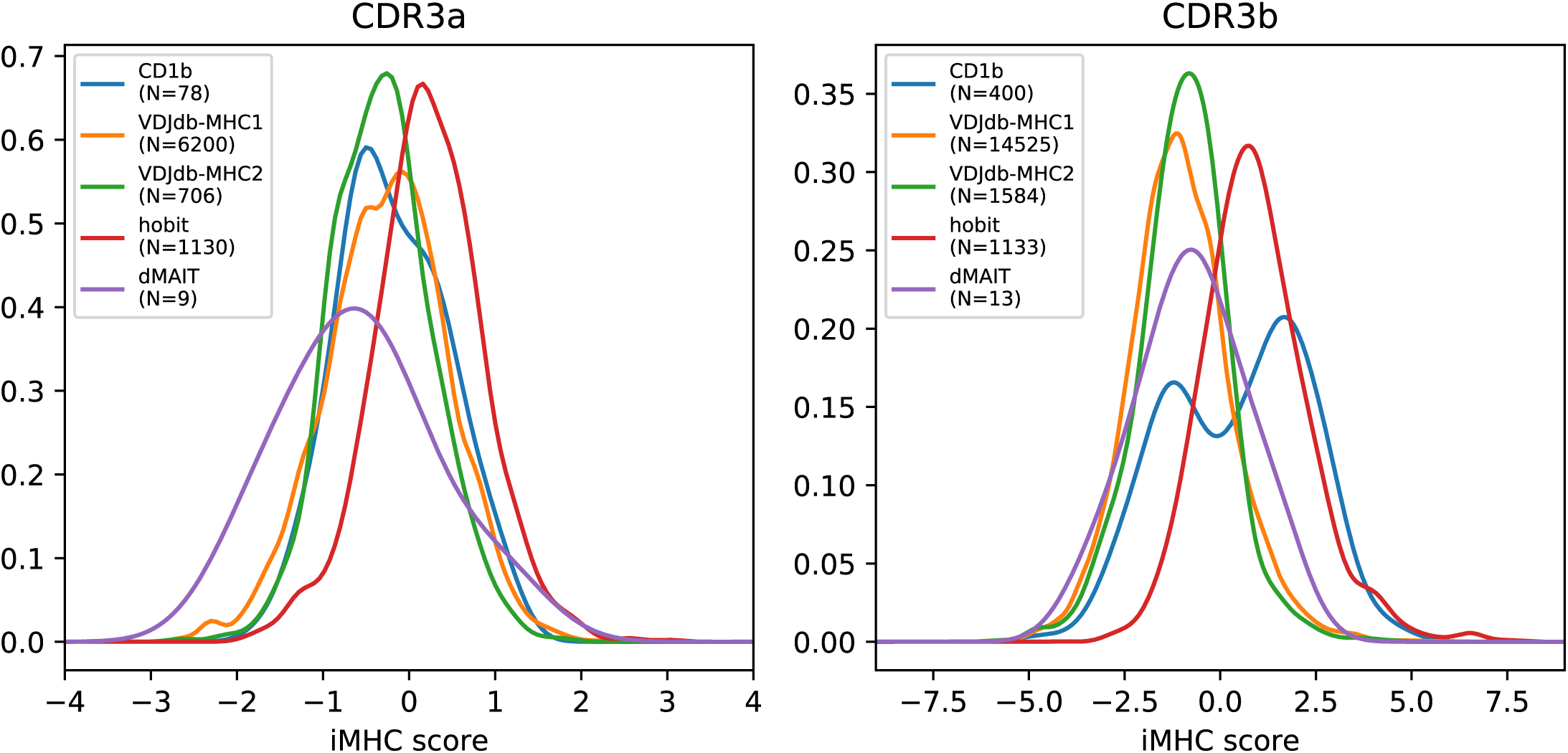
Single-chain iMHC score distributions for TCR subsets. Score distributions for CDR3α repertoires are shown on the left and for CDR3β repertoires on the right. Single-chain variants of the iMHC score were fit with L1-regularized logistic regression just as for the paired iMHC score. Subset labels are as follows: ‘CD1b’, GMM:CD1b-tetramer sorted T cells from DeWitt et al.^44^; ‘VDJdb-MHC1’, TCRs reported to bind to MHC class 1 presented epitopes in the VDJdb database^45^; ‘VDJdb-MHC2’, TCRs reported to bind to MHC class 2 presented epitopes in the VDJdb database; ‘dMAIT’, diverse MAIT TCR sequences from Gherardin et al.^46^ 2016); ‘hobit’, T cells belonging to the *HOBIT+*/*HELIOS+* population in *10x_200k* donors 1, 3, and 4.

**Figure S11:**
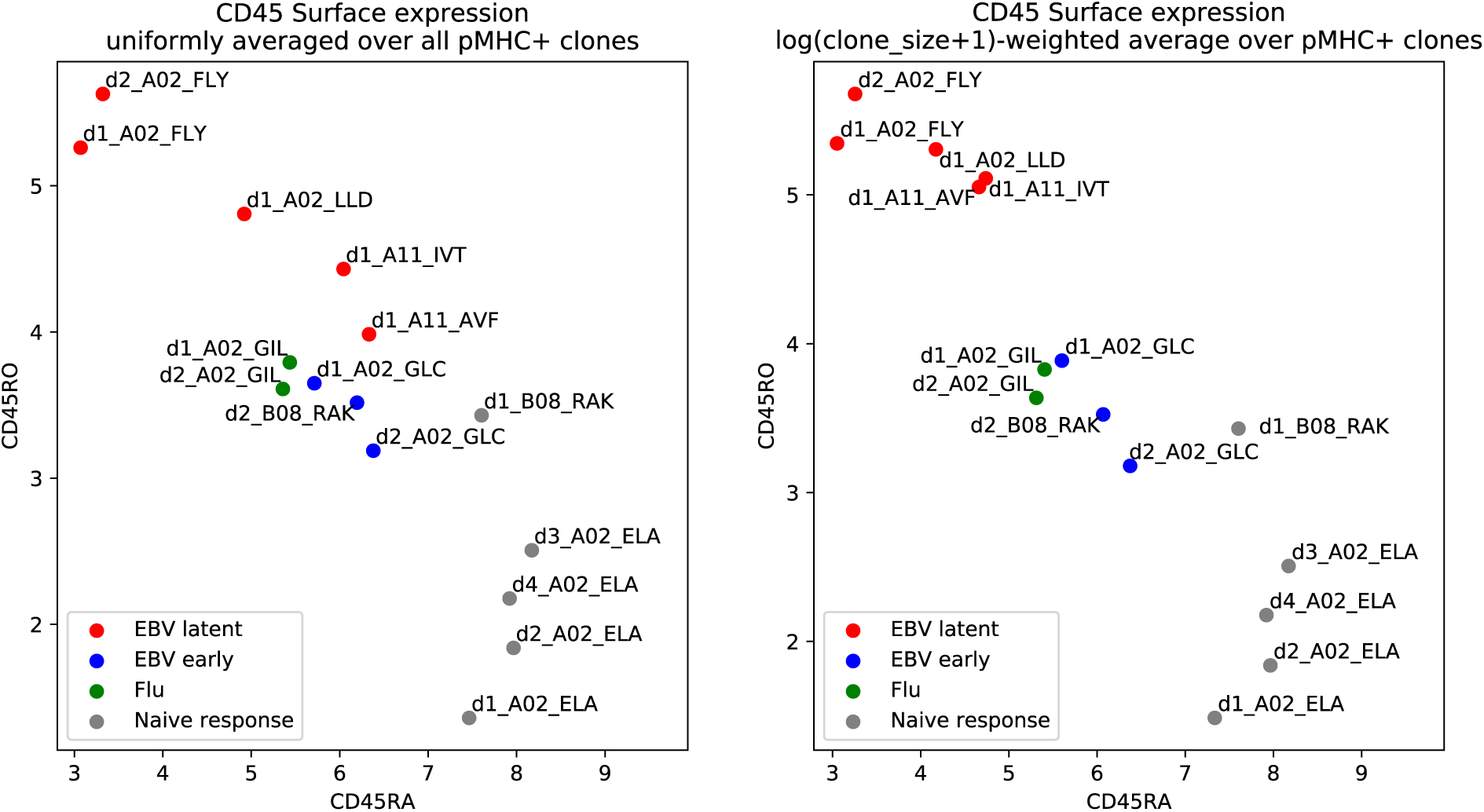
Epitope-specific T cell populations differ in CD45RA/RO expression levels. Log-transformed read counts for DNA-barcoded anti-CD45RA (x-axis) and anti-CD45RO (y-axis) antibodies, averaged over pMHC+ clonotypes, are plotted for the pMHCs shown in main text **Figure 10**. In the panel on the left, clonotypes are weighted equally, while in the panel on the right, larger clonotypes are given more weight (proportional to the logarithm of the clone size) to better reflect the underlying distribution of cells (particularly for the d1_A11 pMHCs, both of which have a relatively large number of positive cells distributed unevenly among a small number of clonotypes).

**Figure S12:**
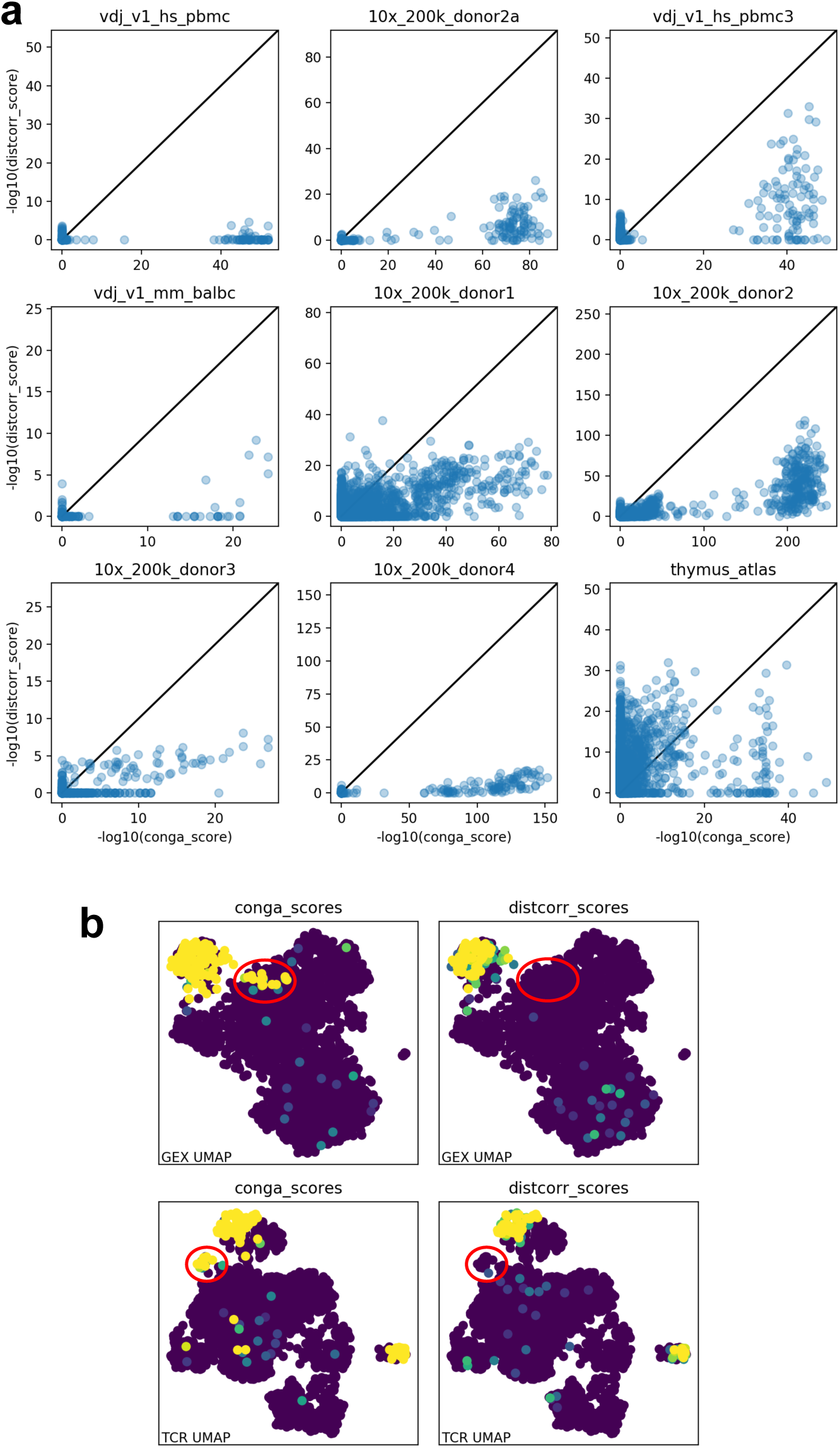
Comparison of graph-based and distance-based measures for assessing GEX/TCR correlation. Comparison of CoNGA scores to a distance-based score (‘distcorr’) that measures, for each clonotype, the degree of correlation between the GEX and TCR distances from that clonotype to all other clonotypes in the dataset. Correlation is assessed using the Pearson correlation coefficient and associated P-value as returned by the scipy.stats.linregress function. **(a)** Scatter plots directly comparing the (negative log_10_-transformed) significance scores assigned to each clonotype in the datasets. The majority of points with significant *P* values lie below the *y* = *x* line, indicating that the CoNGA graph-overlap measure assigns higher significance than distance correlation. One difficulty with raw distance correlation is that it doesn’t discriminate between a set of clonotypes with low distances for both measures (nearby in GEX and in TCR space), on the one hand, and a set of clonotypes with high distances for both measures (far away in GEX and in TCR space), on the other: both increase the correlation coefficient, so a tight cluster in GEX and TCR space (like MAIT cells) can artificially elevate distcorr scores for distant clones. **(b)** CoNGA and distcorr scores mapped to the GEX and TCR landscapes for *10x_200k_donor2a*. The red ellipses indicate the A*02:M1_58_ clones, which appear to be completely missed by the distcorr measure.

